# Hierarchical processing of sensory information across topographically organized thalamocortical-like circuits in the zebrafish brain

**DOI:** 10.1101/2025.09.15.675867

**Authors:** Anh-Tuan Trinh, Anna Maria Ostenrath, Ignacio del Castillo-Berges, Susanne Kraus, Fanchon Cachin, Bram Serneels, Koichi Kawakami, Emre Yaksi

## Abstract

Thalamocortical projections contribute to the spatial organization and functional hierarchies of the mammalian cortex. Primary sensory cortices receive topographically segregated information from first-order thalamic nuclei, which process distinct sensory modalities. In contrast, higher-order cortical regions integrate information from multiple information channels. While such hierarchical processing and integration of information is the foundation for neural computations in the mammalian cortex, the fundamental principles of thalamocortical computations in non-mammalian vertebrates remains unexplored. The zebrafish pallium, located in the dorsal telencephalon, is regarded as the homolog of the mammalian cortex. However, it remains unclear how the zebrafish pallium receives and processes sensory information, and how the architecture and function of these processes compare to the thalamocortical circuits in other vertebrates. Using anatomical tracing, electrophysiological circuit mapping, and *in vivo* Ca^2+^ imaging, we revealed a thalamocortical-like pathway in the zebrafish brain. We found that the preglomerular nucleus (PG) is the primary source of visual and mechano-vibrational information to the zebrafish pallium. We demonstrated that PG neurons and their pallial projections exhibit sensory-specific and topographically organized responses. In contrast, the sensory responses of pallial neurons display multiple layers of topographically organized hierarchies, ranging from simple sensory-specific responses to multimodal and coincidence-detecting nonlinear responses. Notably, we observed a progressive increase in the complexity of sensory computations, which is organized topographically along the posterior-anterior axis of the zebrafish pallium. Collectively, our results suggest that hierarchical sensory processing across topographically organized pallial regions is a conserved functional feature of the vertebrate pallium.

## INTRODUCTION

A fundamental feature of the mammalian cortex is the hierarchical processing of information distributed across the brain (1–10). These cortical hierarchies enable parallel neural computations ranging from sensory encoding (11–14) to the integration of multiple information streams (1, 4, 6, 15). They often utilize multiple coding strategies (16–19) that give rise to cognitive function (9, 16, 20, 21). Strikingly, growing molecular, anatomical, and physiological evidence from birds (22–26), reptiles (27–32), amphibians (33, 34) and fish (35–42) suggests that the vertebrate pallium, an ancestral brain region from which the mammalian cortex evolved, exhibits multiple topographically organized hierarchies. However, revealing whether these pallial hierarchies operate through convergent or divergent computational principles requires the simultaneous monitoring of neural activity across thousands of individual neurons distributed across multiple brain regions (9, 43–45), a major experimental challenge in most vertebrates.

Thalamocortical projections play a fundamental role in shaping the hierarchical organization and specialization of the mammalian cortex (3, 6, 46, 47). Primary sensory cortical areas receive spatially organized inputs from first-order thalamic nuclei, each dedicated to specific sensory modalities (13, 14). For example, the first order visual thalamus, the lateral geniculate nucleus (LGN), receives direct retinal inputs and relays this information to V1 neurons encoding visual orientation, direction, and spatial location (14, 48–50). In contrast, higher-order and associative cortical regions integrate signals from multiple modalities through cortico-thalamic loops mediated by high-order thalamic nuclei (2, 3, 47, 51). For example, a higher order thalamus, the pulvinar nucleus, receives information primarily from the superior colliculus (SC), visual and high-order cortical areas (14, 52). They then relay information to higher order visual, motor and frontal cortices involved in recognition of complex shapes and objects, spatial attention and guiding movements (52, 53). Hence, the first order and high-order thalamic projections to the mammalian cortex represent parallel pathways with complementary functions. Likewise, these two parallel pathways, thalamofugal (e.g. retina->LGN->hyperpallium) and tectofugal (e.g. retina->SC-> nucleus rotundus −>entopallium) are present in reptiles (54–56) as well as birds (50, 56–59) and were shown to be involved in visual motion, visual perception, and visual-spatial processing (27, 58, 60, 61).

Teleost fish, representing half of all vertebrate species, also possess a pallium located in the dorsal telencephalon, composed of spatially organized regions that are both molecularly (36, 42, 62–65) and functionally (66–79) distinct. Genetic ablations and lesions of distinct teleost pallial regions have established a strong link to their role in various adaptive behaviors and learning (66–69, 72, 74). Intriguingly, the teleost thalamus proper (developmentally comparable to the first order thalamus) send very weak projections to the pallium (80, 81). Yet, a parallel teleost sensory relay region, the preglomerular nucleus (PG), was shown to receive tectal innervations while also projecting massively to the pallium. Hence, this suggests that the PG is anatomically comparable to the tectofugal relays in other amniotes like the nucleus rotundus in reptiles and pulvinar nucleus in mammals. Despite the PG’s central role as a main driver of sensory information to the teleost pallium, how sensory information is encoded within PG and subsequently transformed and integrated across hierarchies of pallial circuits remains largely unknown.

Here, we used zebrafish, a small and genetically tractable teleost to investigate how the PG encodes sensory information, how it transmits these inputs to the pallium, and what computational principles and hierarchies govern sensory processing and integration across these pallial circuits. By using anatomical tracing, we showed that PG is the primary input hub to the pallium, channelling information from the diencephalon and mesencephalon. Next, using *in vivo* calcium imaging, we showed that PG neurons and their pallial axon terminals encode and relay visual and vibration information to topographically distinct pallial targets. We also observed that topographically distinct regions of the zebrafish pallium integrate sensory information differently with decreasing selectivity and increasing complexity that is hierarchically organized along the posterior to anterior axis. Our results suggest that the zebrafish pallium has evolved convergent principles of sensory processing hierarchies primarily using the tectofugal rather than thalamofugal pathways.

## RESULTS

### The preglomerular nucleus is the primary source of inputs to the zebrafish pallium

We first asked how the zebrafish pallium receives sensory information, beyond olfaction. To do this we used 21-28 days-old juvenile zebrafish with optically accessible brains. At this age zebrafish are capable of performing cognitively demanding behaviors typically associated with pallial computations, including learning (82–84), social interactions (85–87) and adaptative behaviors (76, 88–90). An earlier study reported the lack of GFP-expressing PG axons in 4-week-old juvenile zebrafish pallium, suggesting that the PG projections to the pallium may develop in older zebrafish (91). To circumvent potential problems with the developmental changes of GFP expression, we performed local iontophoreresis injections of TMR-dextran, a red fluorescent neural tracer (92–94), into the anatomically well-defined PG in juvenile zebrafish whole-brain explants (Fig. 1A-B). We observed anterogradely transported fluorescent dye in the PG axons innervating the pallium of both 2- and 3-week-old juvenile zebrafish (Fig. 1B, fig. S1, fig. S2). PG axon terminals were only present in the ipsilateral pallial hemisphere, predominantly in distinct pallial regions: the dorsal medial (Dm), dorsal central (Dc), and dorsal lateral (Dl) telencephalon (Fig. 1B, Fig. 1C). The same injections also revealed retrograde transport of the fluorescent tracer, labeling cell bodies of PG-innervating neurons across the zebrafish brain. PG-innervating neurons were prominent in the ipsilateral hemisphere of diencephalic (Hyp, Th, PO, TLa) and mesencephalic (OT, SGN, TS) nuclei (Figure 1D). Surprisingly, we also observed a group of pallial Dm neurons projecting back to PG (Fig. 1C).

**Fig. 1.**
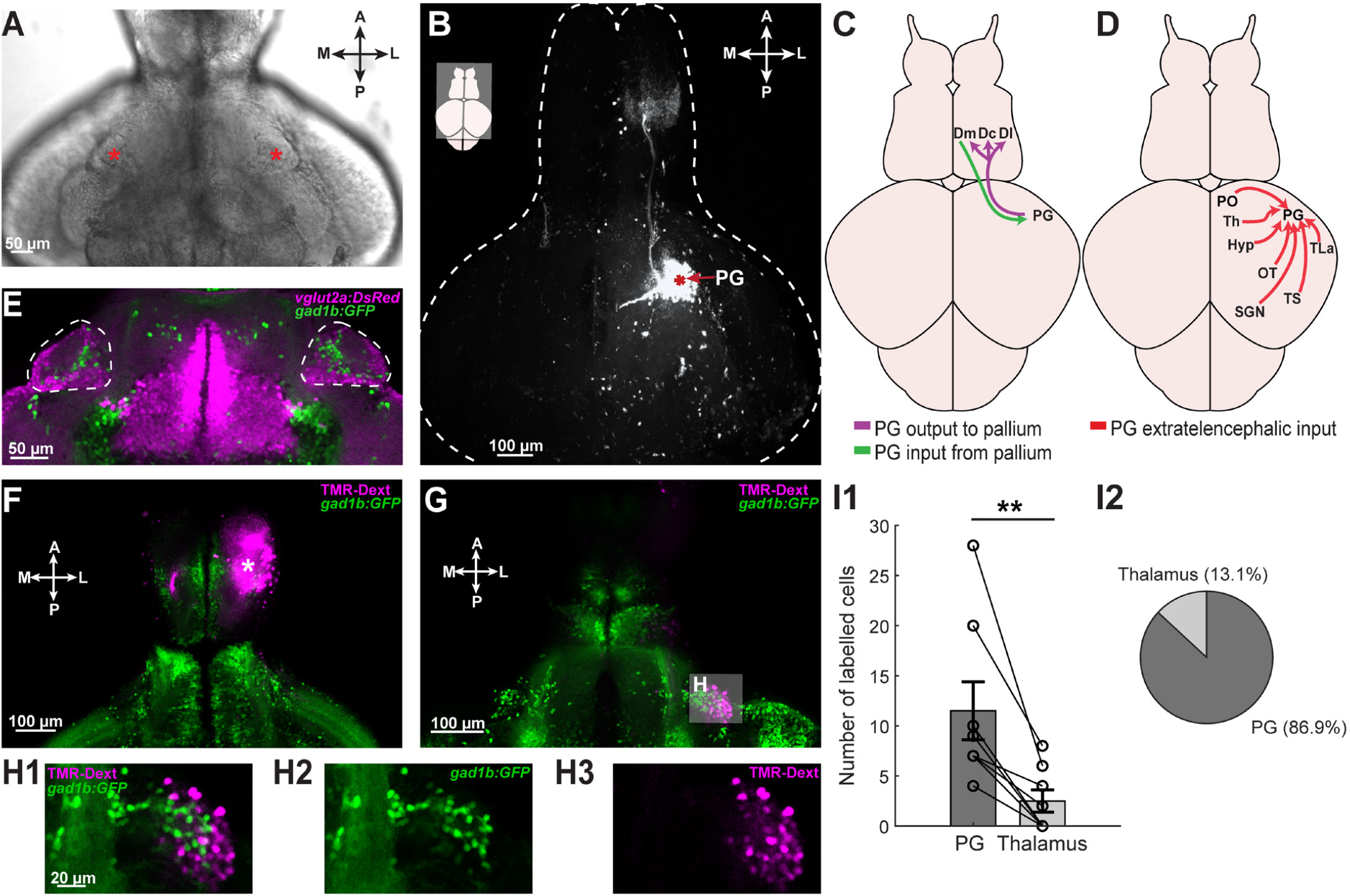
The preglomerular nucleus is the major source of diencephalic inputs to the zebrafish pallium. **A**. Example brightfield image of a dissected juvenile fish brain explant taken from the ventral side. Red stars mark the anatomically distinct lobes of the PG. **B**. Example neurotracer (TMR-Dextran) injection in the PG of a juvenile fish brain explant. Red star marks the injection site. The white dashed line marks the outline of the brain. **C**. Summary scheme of the observed extrinsic connectivity between the PG and the zebrafish dorsal telencephalon/pallium (N = 14 fish). **D**. Summary scheme of the observed connectivity between the PG and diencephalic/mesencephalic structures (N = 14 fish). **E**. Fluorescent image of a dissected juvenile zebrafish brain explant of the *Tg(vglut2a:DsRed;gad1b:GFP)* fishline; vglut2a neurons are in magenta and gad1b neurons in green. The white dashed line highlights the location of the PG. **F-G**. Example neurotracer (TMR-dextran, in magenta) injection in the pallium of a juvenile *Tg(gad1b:GFP)* zebrafish (N = 8 fish). The dorsal plane is shown in F, while the ventral plane is shown in G. **H1-3**. Magnified view of the ipsilateral PG (in G) following the neurotracer injection in the pallium in F. **I1**. Number of retrogradely labelled neurons within the PG and thalamus proper following neurotracer injections in the pallium (N = 8 fish, ** p < 0.01, Wilcoxon signed-rank test). **I2**. Pie chart illustrating the mean fraction of labelled neurons within the PG and thalamus proper following pallial neurotracer injections (N = 8 fish). Abbreviations: Dm: dorsal medial telencephalon, Dc: dorsal central telencephalon, Dl: dorsal lateral telencephalon, PG: preglomerulus nucleus, PO: pre-optic area, Th: thalamus proper, Hyp: hypothalamus, TLa: lateral torus, OT: optic tectum, TS: torus semicircularis, SGN: secondary gustatory nucleus. A: anterior, M: medial, L: lateral, P: posterior.

The mammalian thalamus is primarily composed of cortical projecting glutamatergic neurons (14, 95). Yet, GABAergic thalamic interneurons and reticular neurons do not project to the cortex. We observed that the zebrafish PG is also composed of both glutamatergic (80.34 ± 1.37% *Tg(vglut2a:DsRed)* (96) labelled) and GABAergic (19.66 ± 1.37%, *Tg(gad1b:GFP)* (97) labelled) neurons (Fig. 1E). To investigate which population of PG neurons project to the pallium, we injected TMR-dextran broadly into the pallium (Fig. 1F) and examined the retrogradely labelled cells in the zebrafish diencephalon (Fig. 1G). These experiments yielded a large fraction of retrogradely labelled PG neurons only in the ipsilateral hemisphere (magenta, Fig. 1G-H). None of these pallial-projecting PG neurons were co-labelled with gad1b (green, Fig. 1H). Additionally, these retrograde labelling experiments also revealed that PG is the primary input region of the zebrafish pallium, with only a small fraction (~13%) of pallial-projecting retrogradely labelled neurons located in the zebrafish thalamus proper (fig. S3A). Finally, we asked whether neurons in distinct PG subnuclei may have preferential projections to distinct pallial regions. To answer this, we performed spatially restricted injections of TMR-dextran into distinct pallial regions Dl (fig. S3B) and Dm (fig. S3C). We observed that lateral PG neurons are retrogradely labelled upon Dl micro-injections (fig. S3D), and medial PG neurons are labelled upon Dm micro-injections (fig. S3E). Altogether, our results suggest that PG is the primary glutamatergic hub relaying diencephalic and mesencephalic information to distinct zebrafish pallial regions through anatomically distinct PG sub-regions.

### The zebrafish preglomerular nucleus is composed of functionally heterogeneous subnuclei

Different mammalian thalamic subnuclei exhibit varying levels of spontaneous ongoing activity with distinct features (98, 99). To examine whether the PG is composed of multiple functional subnuclei, we measured the ongoing neural activity of the entire PG *in vivo* using two-photon calcium imaging in head-restrained *Tg(elavl3:H2B-GCaMP6s)* (100) juvenile zebrafish at 10-14 dpf (Fig. 2A). To characterize the spatio-temporal features of ongoing PG activity, we grouped ensembles of PG neurons with similar spontaneous calcium dynamics using k-means clustering (70, 76, 90, 101) (Fig. 2B). We identified that PG activity can be represented with ~6 clusters (fig. S4A-B). We observed that functional clusters/ensembles of PG neurons with similar calcium dynamics are topographically organized into distinct PG zones (Fig. 2C). To quantify this functional topography further, we plotted the average pairwise correlation of PG neurons as a function of distance between pairs (70, 71, 76, 90, 101–103). We observed that nearby PG neurons exhibit higher correlations compared to distant neurons (Fig. 2D, fig. S4C). Next, we assessed the stability of the PG ensembles by quantifying the likelihood of PG neuron pairs to remain in the same cluster during two consecutive time periods, a measure we termed “cluster fidelity” (70, 71, 76, 90, 101, 102). This analysis revealed that a significantly larger fraction of PG neuron pairs remained in the same cluster, above chance level (Fig. 2E). In line with this, we observed that correlations between PG neuron pairs remained stable during consecutive time periods (Fig. 2F). Our results demonstrate that the zebrafish PG is composed of functionally heterogeneous and topographically organized subnuclei.

**Fig. 2.**
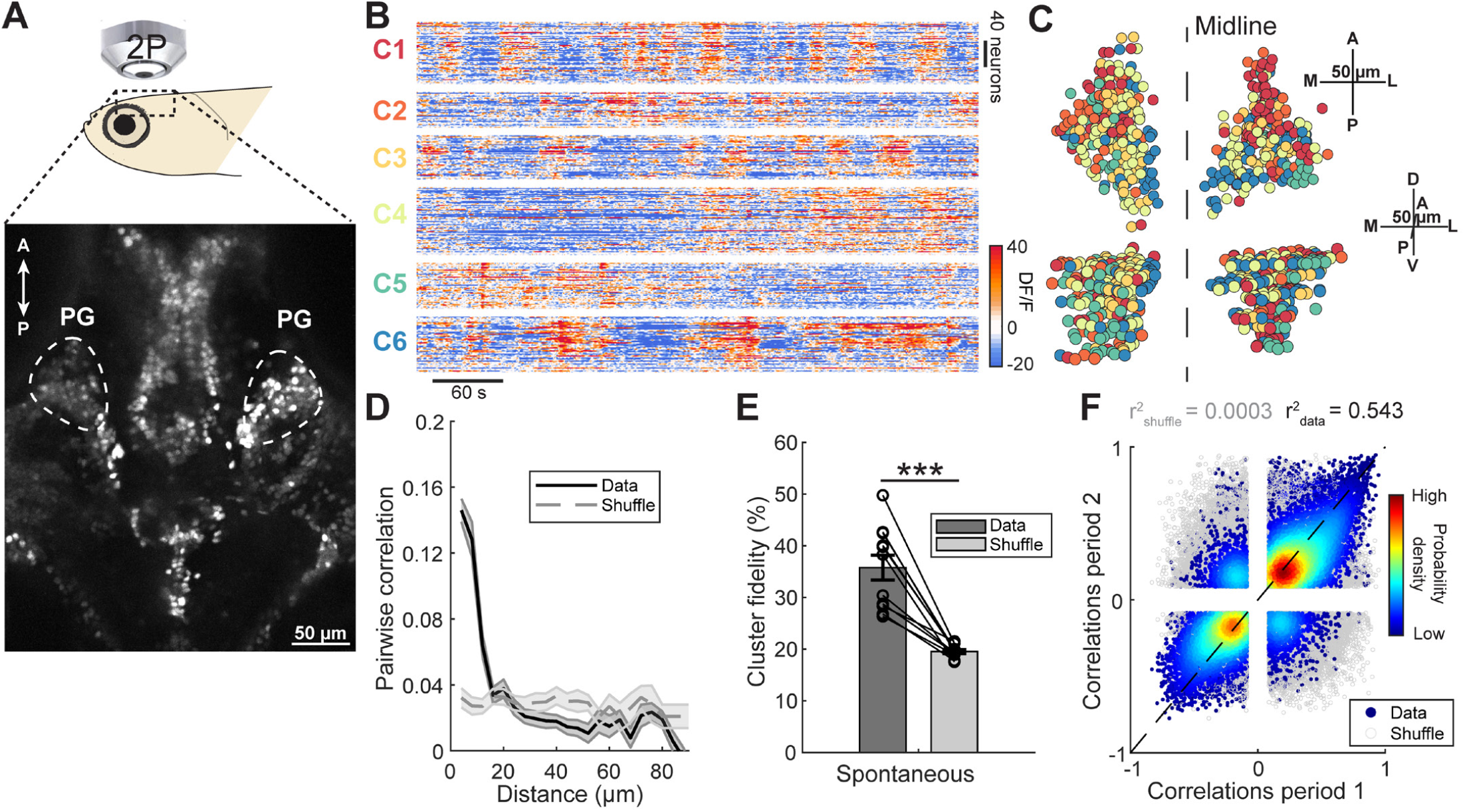
Preglomerular nucleus neurons form topographically organized functional ensembles based on their spontaneous activity. **A**. Example two-photon microscopy image of a *Tg(elavl3:H2B-GCaMP6s)* zebrafish, *in vivo*. White dashed lines mark the borders of PG. **B**. Example in vivo spontaneous ongoing activity traces of PG neurons clustered using k-means. Warm colors represent increased neural activity **C**. 3D reconstruction of PG neurons in the same example fish. The neurons are color-coded by the cluster identity, marked in B. **D**. Pairwise correlations of the PG neurons’ spontaneous ongoing activity as a function of the distance between neuron pairs. The grey dashed line represents pairwise correlations when the distances are shuffled. The shaded region denotes the standard error of the mean (SEM) (N = 11 fish). **E**. Cluster fidelity of the PG neurons during two consecutive spontaneous activity periods, in comparison to a shuffled distribution of the cluster identities. **F**. Pairwise Pearson’s correlations of PG neuron pairs during two consecutive periods of spontaneous activity. Only significant correlations (p > 0.05) are plotted. The probability density of pairwise correlations in the scatter plot is represented with a color gradient. The grey dots represent the pairwise correlations obtained from a shuffled distribution. The correlation coefficient for the shuffled (in gray, r^2^ = 0.0003) and observed data (in black, r^2^ = 0.543) were obtained following a linear fit represented by the black dashed line. The Error bars denote the SEM (N = 11 fish, *** p < 0.001, paired Wilcoxon signed-rank test).

### The preglomerular nucleus exhibits selective and topographically organized responses to different sensory modalities

In amniotes, the thalamus is divided into distinct first-order subnuclei that encode information from distinct sensory modalities (1, 13, 14, 95). To investigate how the zebrafish PG encodes sensory information, we delivered series of red light flashes and mechanical vibrations in separate trials while measuring the PG neuronal activity *in vivo* in *Tg(elavl3:H2B-GCaMP6s)* (100) juvenile zebrafish. We observed that a fraction of PG neurons was excited by light and vibrations (Fig. 3A1, 3B1, in red), while another fraction was inhibited (Fig. 3A2, 3B2, in blue). Next, we asked whether PG neurons are selective for individual sensory modalities. We plotted the distribution of response amplitudes of those PG neurons with significant excitatory responses. Most PG neurons responded to either light or vibration stimuli (Fig. 3C). We then plotted the proportion of PG neurons that responded exclusively to either light or vibration stimuli, as well as those that responded to both, which we termed multi-sensory neurons (Fig. 3D). We observed that a significantly larger fraction of PG neurons exclusively respond to one sensory modality compared to multi-sensory neurons, highlighting the sensory selective nature of the PG neurons.

**Fig. 3.**
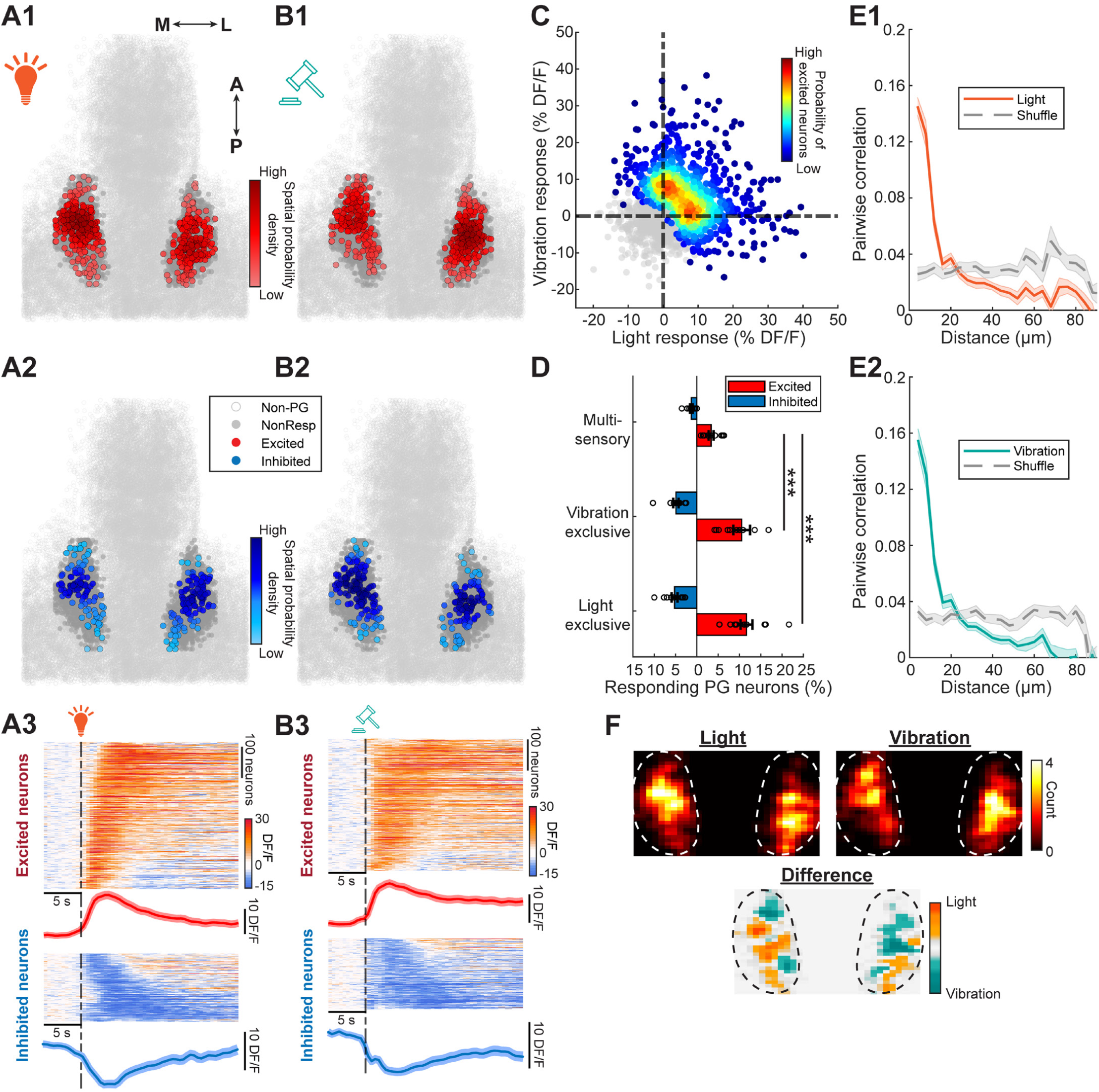
Preglomerular nucleus neurons exhibit selective sensory responses. **A-B**. Overlaid 2D reconstruction of excited (in red, A1-B1) and inhibited (in blue, A2-B2) PG neurons from all fish, in response to light (A) and vibration (B) stimulation, in vivo. The color gradient represents the spatial probability density of the responding PG neurons. Non-responding PG neurons are in dark gray while neurons outside of PG are in light gray. Time courses of the PG neurons’ calcium signals in response to light (A3) and vibrations (B3). Warm colors indicate increased neural activity; cold colors indicate decreased activity. Mean calcium signals are shown at the bottom of each heatmap, shades indicate the standard error of the mean (N = 11 fish). **C**. Mean responses of all individual PG neurons upon light and vibration stimulation. The probability density of PG neurons with significant excitation upon light or vibration stimulation is color-coded. Non-excited neurons are shown as gray dots. **D**. The fraction of responding PG neurons excited (red) and inhibited (blue) by either light or vibration (exclusive), or by both stimuli (multi-sensory), in individual fish (black open circles). **E**. The pairwise correlation of the PG neurons’ responses to light (E1) and vibration (E2) stimulation as a function of the distance between neuron pairs. The grey dashed line represents the pairwise correlations when the distances are shuffled. The shaded region denotes the standard error of the mean. (N = 11 fish). **F**. Top row: The locations of all light (left) and vibration (right) excited PG neurons from all fish are spatially aligned and overlayed in a 2D histogram. Warmer colors denote higher neuron counts per bin. The difference between the two 2D histograms is shown at the bottom. Warm color highlights a spatial preference for the light excited PG neurons while green colors denote a preference for the vibration excited PG neurons (N = 11 fish). The dashed line marks the anatomical boundaries of the PG. Error bars are standard error of the mean (N = 11 fish, *** p < 0.001, Wilcoxon rank-sum test).

Next, inspired by the topographically organized spontaneous ongoing PG activity (Fig. 2), we asked whether distinct subregions of PG exhibit topographically organized sensory responses. To quantify the topography of sensory-evoked PG activity, we calculated pairwise correlations of PG neuron responses as a function of distance between them. This analysis revealed significantly higher response correlations between nearby PG neurons, exceeding chance levels (Fig. 3E). We also showed that PG responses were not randomly distributed but exhibited a higher degree of focality above chance levels (fig. S4D). In fact, we observed that neurons within topographically organized PG ensembles (Fig. 2) also responded to sensory stimulation in similar ways (fig. S4E-F), as quantified by a higher than chance cluster fidelity index (fig. S4G). Finally, we examined how PG neurons responding to light and vibrations were spatially distributed in PG by spatially aligning the excited sensory-responding neurons from all PG recordings. At first, the two-dimensional spatial histograms of the PG responses looked similar for both light and vibration responses (Fig. 3F, top). However, subtracting these spatial histograms revealed that the posterior-lateral PG neurons have a higher preference for light (orange) while the rostral-medial PG neurons prefer the vibration stimuli (teal, Fig. 3F, bottom). Altogether, these results revealed that PG neurons exhibit selective sensory responses, and partially overlapping but segregated zones of PG prefer different sensory stimuli.

### Preglomerular nucleus axons innervating zebrafish pallium are organized into distinct topographical zones selective for different sensory modalities

Axonal projections of relay neurons from distinct thalamic nuclei play an important role in the functional and topographical organization of the amniote cortex (2, 14), but whether similar principles are also observed in teleost fish is unclear. To examine this, we used a transgenic fishline *Tg(gSAIzGFFD707A:Gal4;UAS:GCaMP6s)* expressing GCaMP6s in the PG neurons and their axonal projections to the zebrafish pallium. We confirmed that the GCaMP6s-expressing axons in the pallium originate from the PG by injecting TMR-dextran into the PG, which showed strong overlap with the GCaMP6s and TMR-dextran signals (Fig. 4A). Given our observation of structured spontaneous ongoing PG activity (Fig. 2), we first investigated the ongoing PG axonal calcium signals in the zebrafish pallium (Fig. 4B). We grouped ensembles of PG axonal pixels with similar ongoing activity using k-means clustering (70, 71, 76, 90, 101) (Fig. 4C, fig. S5A-B) and observed that the PG axons in distinct clusters were topographically organized (Fig. 4D). In fact, nearby PG axonal pixels exhibited more positively correlated calcium dynamics (Fig. 4E, fig. S5C). Ensembles of PG axons (Fig. 4F) and correlations between PG axonal calcium dynamics also remained stable across different time periods (Fig. 4G). These results revealed that the spontaneous activity of the pallial PG axonal terminals in zebrafish are topographically organized.

**Fig. 4.**
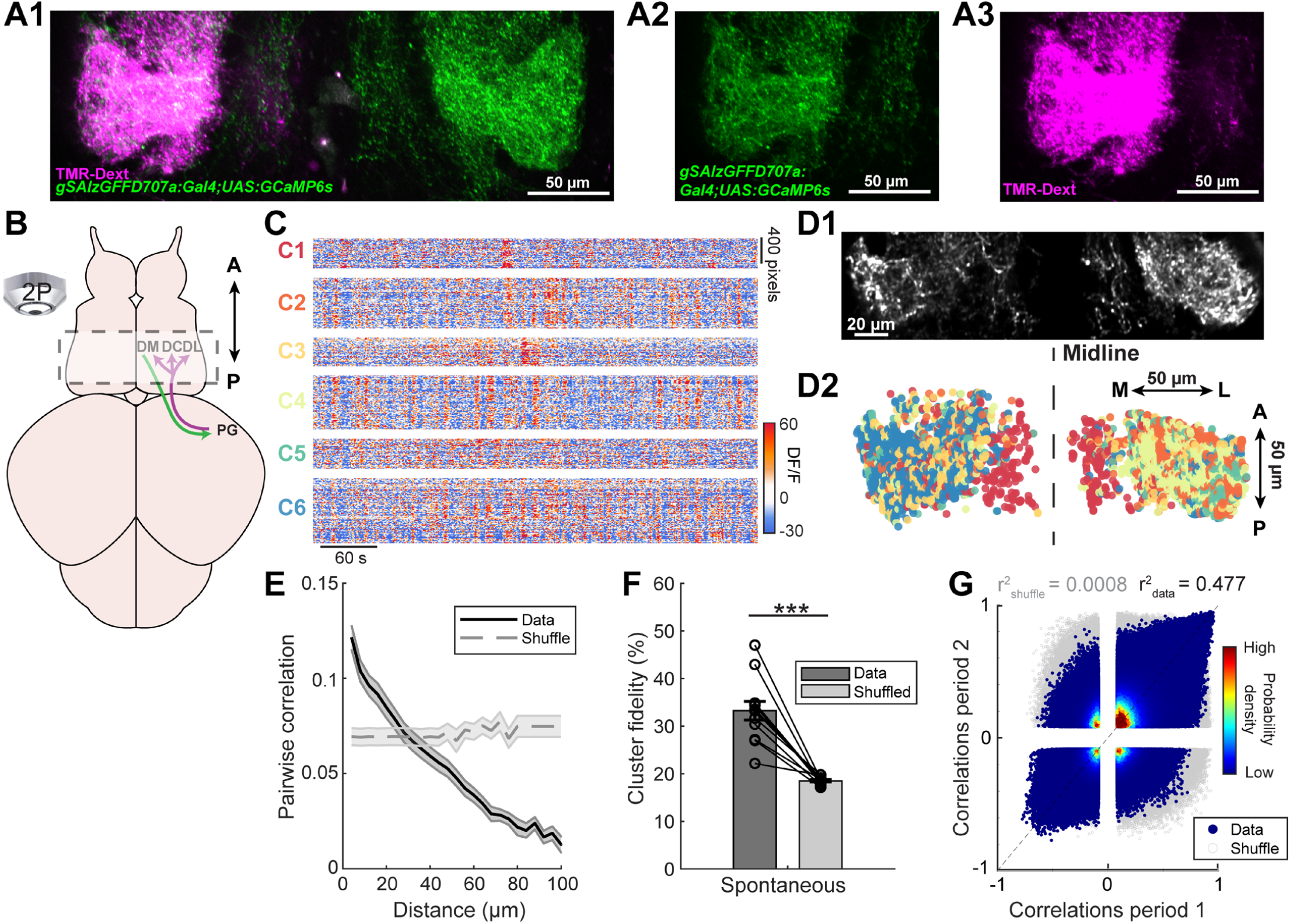
The axons of preglomerular nucleus neurons form topographically organized ensembles in the zebrafish pallium. **A**. Two-photon microscopy image of the PG axons in the zebrafish pallium expressing the transgenic calcium indicator GCaMP6s (green) in *Tg(gSAlzGFFD707A:Gal4;UAS:GCaMP6s)* juvenile fish. These GCaMP6s-labelled axons are confirmed to originate from the PG, by co-labeling with TMR-dextran injection in the PG, as in Figure 1B (N = 3 fish). **B**. Scheme illustrating the imaging area and location of the PG axonal projections in the zebrafish pallium, indicated by dashed lines. **C**. Example calcium signals from pallial PG axons (detected in pixel units) during spontaneous ongoing activity, in an individual fish. The axonal activity was clustered using k-means into 6 different clusters. **D1**. Two-photon microscopy image of the PG axons in the same fish as in C. **D2**. 2D reconstruction of the detected PG axons color-coded by the k-means cluster identity from C. **E**. Pairwise correlations of the PG axons’ spontaneous activity as a function of the distance between pixel pairs. The grey dashed line represents the pairwise correlations when the distances are shuffled. The shaded region denotes the standard error of the mean (SEM). (N = 12 fish) **F**. Cluster fidelity of the PG axons during two consecutive spontaneous activity periods, in comparison to a shuffled distribution of the cluster. **G**. Pairwise Pearson’s correlations of the PG axonal pixels during two consecutive periods of spontaneous activity. Only significant correlations (p > 0.05) are plotted. The probability density of the pairwise correlations in the scatter plot is represented with a color gradient. The grey dots represent the pairwise correlations obtained from a shuffled distribution. The correlation coefficient for the shuffled (in gray, r^2^ = 0. 0008) and observed data (in black, r^2^ = 0.477) were obtained following a linear fit, indicated by the dashed line). The error bars are the SEM (N = 12 fish, ***p < 0.001, paired Wilcoxon signed-rank test).

We next examined what sensory information is conveyed to the zebrafish pallium via the PG axons. To do so, we presented the head-restrained juvenile fish with the light and vibration stimuli, while recording the calcium activity of PG axons in the pallium. We observed both excitation and inhibition in the pallial PG axons upon light and vibration stimulation (Fig. 5A-B). We then asked whether these PG axons, like the PG neurons, are selective for individual sensory modalities. The distribution of response amplitudes of these PG axons revealed a strong preference for either light or vibration stimuli (Fig. 5C). Consequently, we observed a significantly larger fraction of PG axon terminals exhibiting significant responses exclusively to either light or vibrations, when compared to a small fraction of multi-sensory axons (Fig. 5D).

**Fig. 5.**
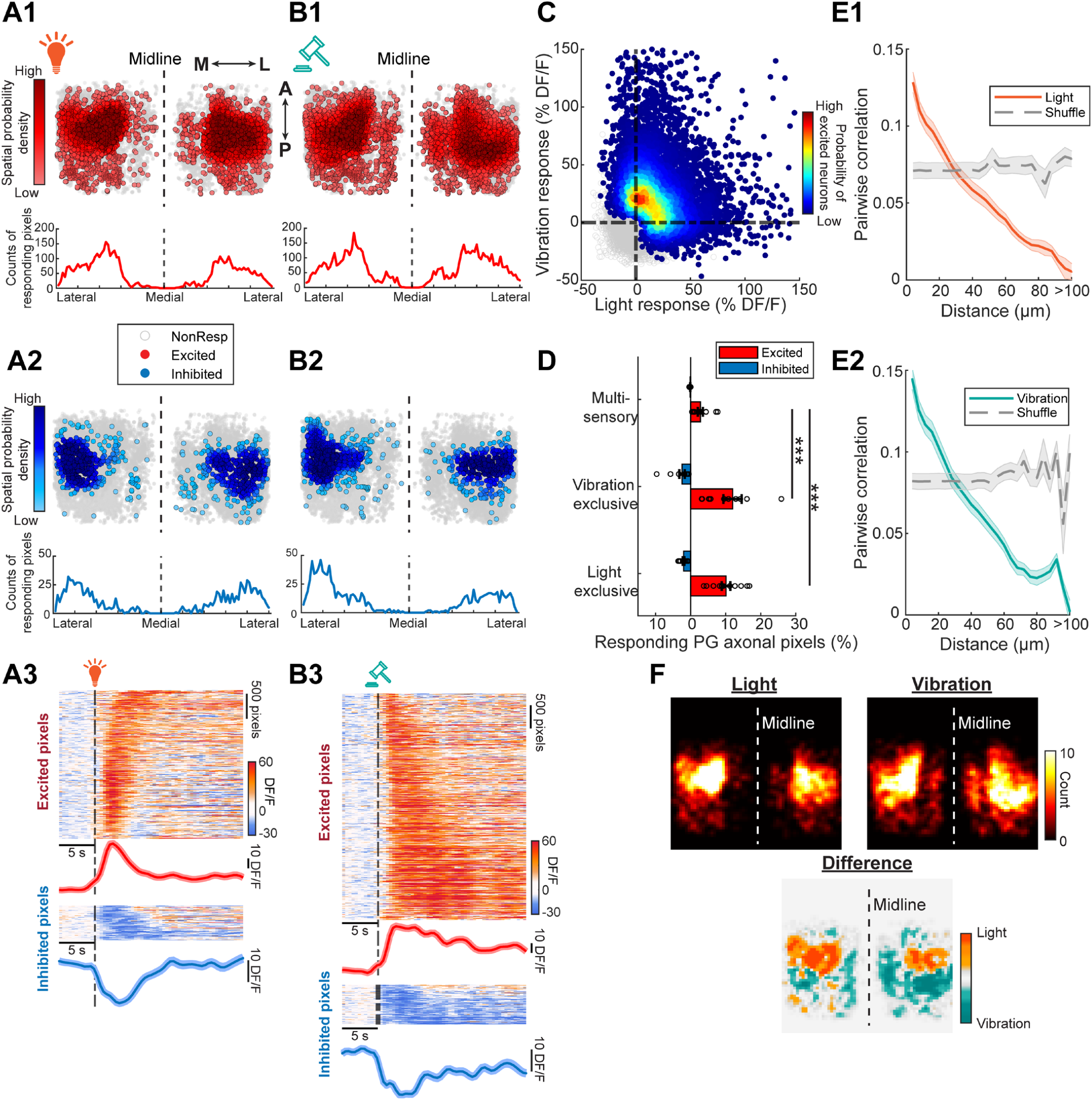
The preglomerular nucleus axons innervating the zebrafish pallium exhibit selective sensory responses. **A-B**. Overlaid 2D reconstruction of the excited (in red, A1-B1) and inhibited (in blue, A2-B2) PG axonal pixels from all fish, in response to light (A) and vibration (B) stimulation, in vivo. The color gradient represents the spatial probability density of responding PG axons. Non-responding PG axons are in gray (N = 12 fish). Time courses of the PG axonal calcium signals in response to light (A3) and vibrations (B3). Warm colors indicate increased neural activity; cold colors indicate decreased activity. Mean calcium signals are shown at the bottom of each heatmap, and the shades indicate the standard error of the mean (N = 8 fish). **C**. Mean responses of all individual PG axonal pixels upon light and vibration stimulation. The probability density of PG axonal pixels with significant excitation upon light or vibration stimulation is color-coded. Non-excited axons are shown as gray dots. **D**. The fraction of responding PG axonal pixels excited (red) and inhibited (blue) by either light or vibration (exclusive), or by both stimuli (multi-sensory), in individual fish. **E**. The pairwise correlation of the PG axonal responses to light (E1) and vibration (E2) stimulation as a function of the distance between axonal pixel pairs. The grey dashed line represents pairwise correlations when the distances are shuffled. The shaded region denotes the standard error of the mean. (N = 12 fish). **F**. Top row: The locations of all light (left) and vibration (right) excited PG axons from all fish are spatially aligned and overlayed in a 2D histogram. Warmer colors denote higher neuron counts per bin. The difference between the two 2D histograms is shown at the bottom. Warm colors highlight a spatial preference for the light excited PG axons while green colors denote a preference for the vibration excited PG axons (N = 12 fish). Error bars represent the standard error of the mean (N = 12 fish, *** p < 0.001, Wilcoxon rank-sum test).

Subsequently, we investigated the topographical organization of sensory responses in PG axons in the pallium and observed that nearby PG axons exhibited more correlated sensory responses (Fig. 5E), and greater focality than chance levels (fig. S5D). Interestingly those PG axons within the same ongoing activity clusters also respond to sensory stimuli, similarly, remaining in the same functional k-means clusters (fig. S5E-G). Finally, we observed that PG axonal terminals that were spatially aligned across all fish, exhibited topographically distinct sensory responses. Specifically, anterior-lateral axons exhibited greater light preferences, and medial and posterior-lateral zones exhibited greater vibration preferences (Fig. 5F). Altogether, our findings revealed that PG axons showed selective sensory responses, segregated to topographically distinct regions of the pallium, similar to the projections from the first order thalamic nuclei onto the primary sensory regions of the mammalian cortex (1, 6, 13, 14).

### Micro-stimulation of the preglomerular nucleus activates topographically distinct pallial regions

Our results showed that PG neurons are the primary diencephalic input to the zebrafish pallium, and they selectively encode different sensory modalities. But how and where in the zebrafish pallium is the information from the PG received? To investigate this, we delivered 2 ms long electrical stimulation into the anatomically well-defined PG in a *Tg(elavl3:H2B-GCaMP6s)* (100) juvenile zebrafish brain explant, using a bipolar glass micro-electrode (70, 93) (Fig. 6A). Upon brief PG micro-stimulations, we observed both excited and inhibited neurons in distinct pallial regions (Fig. 6B). Although PG neurons innervate only the ipsilateral pallium anatomically (Fig. 1B), both ipsi- and contralateral pallium were activated upon PG micro-stimulation. Excitatory responses were significantly more prominent in the ipsilateral pallium, but the distribution of inhibitory responses was similar across both pallial hemispheres (Fig. 6C). The excited neurons were more prevalent in the medial, central, lateral and anterior regions of the pallium, Dm, Dc, Dl, Da (Fig. 6B1-D1). Instead, inhibited neurons were primarily located in the anterior pallium, Da (Fig. 6B2-D2). Micro-stimulating just outside the PG elicited no significant responses in the pallium (fig. S6A-F). Likewise, blocking synaptic connectivity by the bath application of glutamatergic receptor antagonists NBQX/APV (10μM/50μM) wiped-out pallial activation upon PG micro-stimulation (fig. S6G-J). We also tested the reliability of the pallial responses upon PG activation independent of excitation or inhibition by calculating the trial-to-trial Pearsons’s correlations of pallial responses upon PG micro-stimulations (Fig. 6E). We observed that the ipsilateral pallium responded with higher tiral to trial corrrelations and hence exhibited more reliable responses to PG stimulation (Fig.6F-G). Those pallial regions with prominent responses were also more reliably responding (Fig. 6H). Altogether, these results revealed that PG micro-stimulations elicited prominent and topographically organized responses across distinct pallial regions, except the dorsal posterior pallium (Dp).

**Fig. 6.**
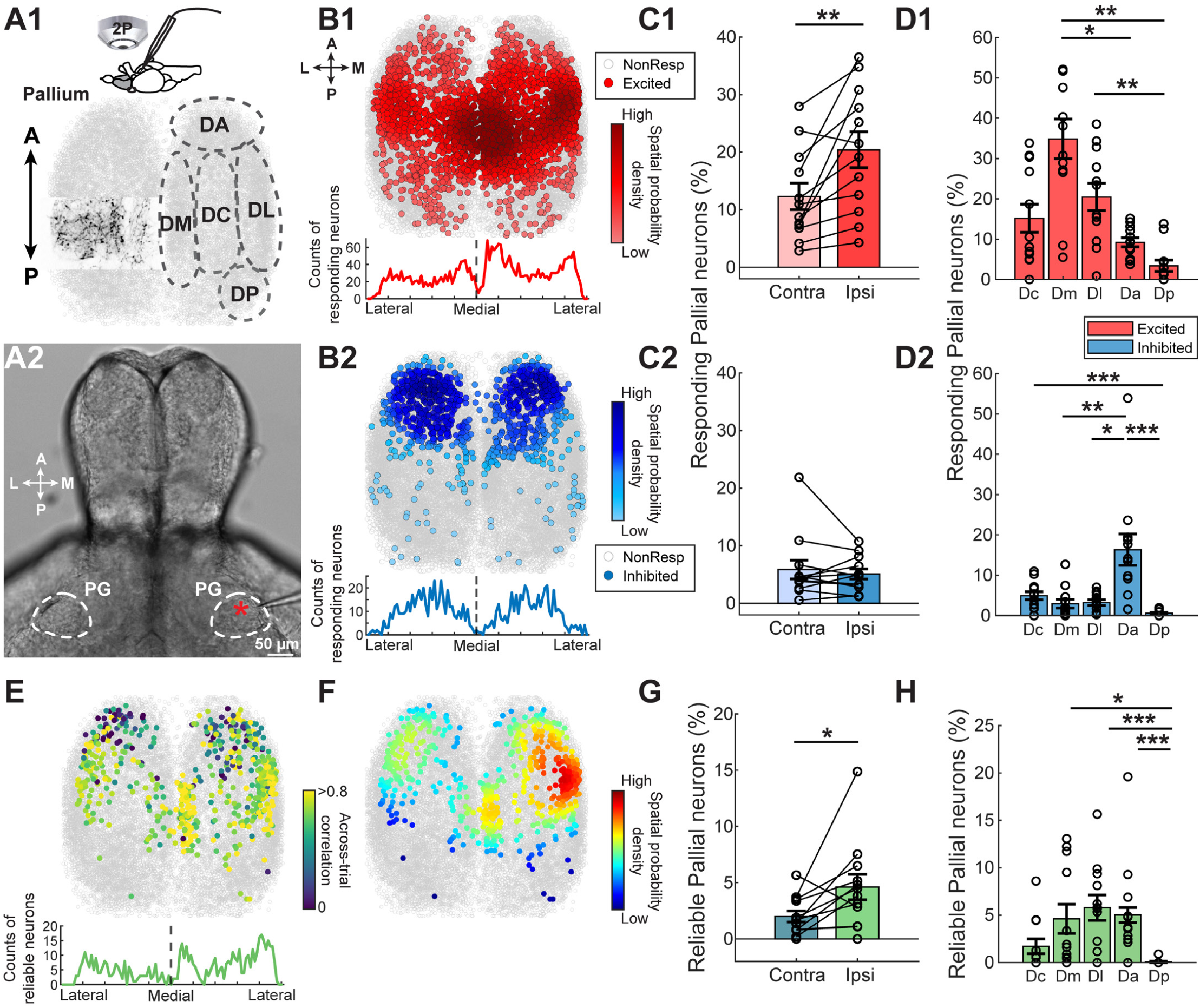
Stimulation of the preglomerular nucleus evokes excitatory and inhibitory responses in anatomically defined pallial regions. **A1**. Top: Scheme representing a micro-electrode stimulation of the PG in juvenile zebrafish brain explants during two-photon calcium imaging of the pallial neurons (N = 12 fish). Bottom: Scheme of the juvenile zebrafish pallium, overlaid with the termination zone of the PG axons (left) and the anatomically identified locations of the pallial regions delineated by the dashed lines (right). **A2**. Brightfield image indicating the location of the microsimulation electrode in the brain explant (denoted by the red star). The PG boundaries are highlighted by white dashed circles. **B1-B2**. Top: 2D reconstruction of all excited (B1, red) and inhibited (B2, blue) pallial neurons and non-responding neurons (in gray) upon PG stimulation. The data from all fish are spatially aligned and overlaid. The color gradient represents the spatial probability density of the responding pallial neurons. Bottom: Number counts of excited (red) and inhibited (blue) neurons along the lateral-medial axis (N = 12 fish). **C**. Fraction of excited (C1) and inhibited (C2) pallial neurons on the ipsi- and contra-lateral hemispheres in reference to the stimulated PG (N = 12 fish, **p < 0.01, paired Wilcoxon signed-rank test). **D**. The fraction of excited (in red, D1) and inhibited pallial neurons (in blue, D2) in anatomically identified pallial regions (N = 12 fish). **E**. Top: 2D reconstruction of the reliably responding pallial neurons upon repeated micro-stimulation trials. Pearson correlations of the trial-to-trial responses of the pallial neurons are color-coded only if the correlation p-value < 0.1. Bottom: Number counts of the reliable responding pallial neurons along the lateral-medial axis (N = 12 fish). **F**. 2D reconstruction of the reliably activated pallial neurons and non-responding neurons (in gray) following the PG stimulation. The color gradient represents the spatial probability density of reliably activated pallial neurons as defined in E. **G**. Fraction of reliably activated neurons across trials plotted per hemisphere (N = 12 fish, *p < 0.05, paired Wilcoxon signed-rank test). The spatial probability density of pallial neurons with is color-coded. **H**. Fraction of reliably activated neurons per delineated brain region (N = 12 fish). All error bars denote the standard error of the mean. (*p-value< 0.05, **p-value < 0.01 & ***p-value < 0.001, Kruskal-Wallis’s test followed by Dunn’s test).

### Sensory responses in the zebrafish pallium are less selective and topographically organized

Having identified the pallial regions receiving PG inputs, we next asked how sensory information is further processed in the zebrafish pallium. To measure sensory responses *in vivo*, we used head-restrained (70, 71, 76, 90, 101, 104) juvenile zebrafish *Tg(eval3:GCaMP6s)* (100), expressing GCaMP6s in all neurons. Upon light and vibration stimulation we observed both excitation and inhibition across the zebrafish pallium (Fig. 7A-B). Unlike the PG neurons and pallial PG axons, a large fraction of pallial neurons exhibited significant multi-sensory responses to both light and vibration stimuli, in addition to those pallial neurons exclusively responding to either light or vibration stimuli (Fig. 7C-D). Nearby pallial neurons exhibited correlated activity (Fig. 7E) and higher focality compared to chance (fig. S7A) upon light and vibration stimulation. Despite a large fraction of multi-sensory pallial neurons, we observed that the lateral (Dl) and medial pallium (Dm), displayed biased preferences for light and vibration stimuli respectively (Figure 7F).

**Fig. 7.**
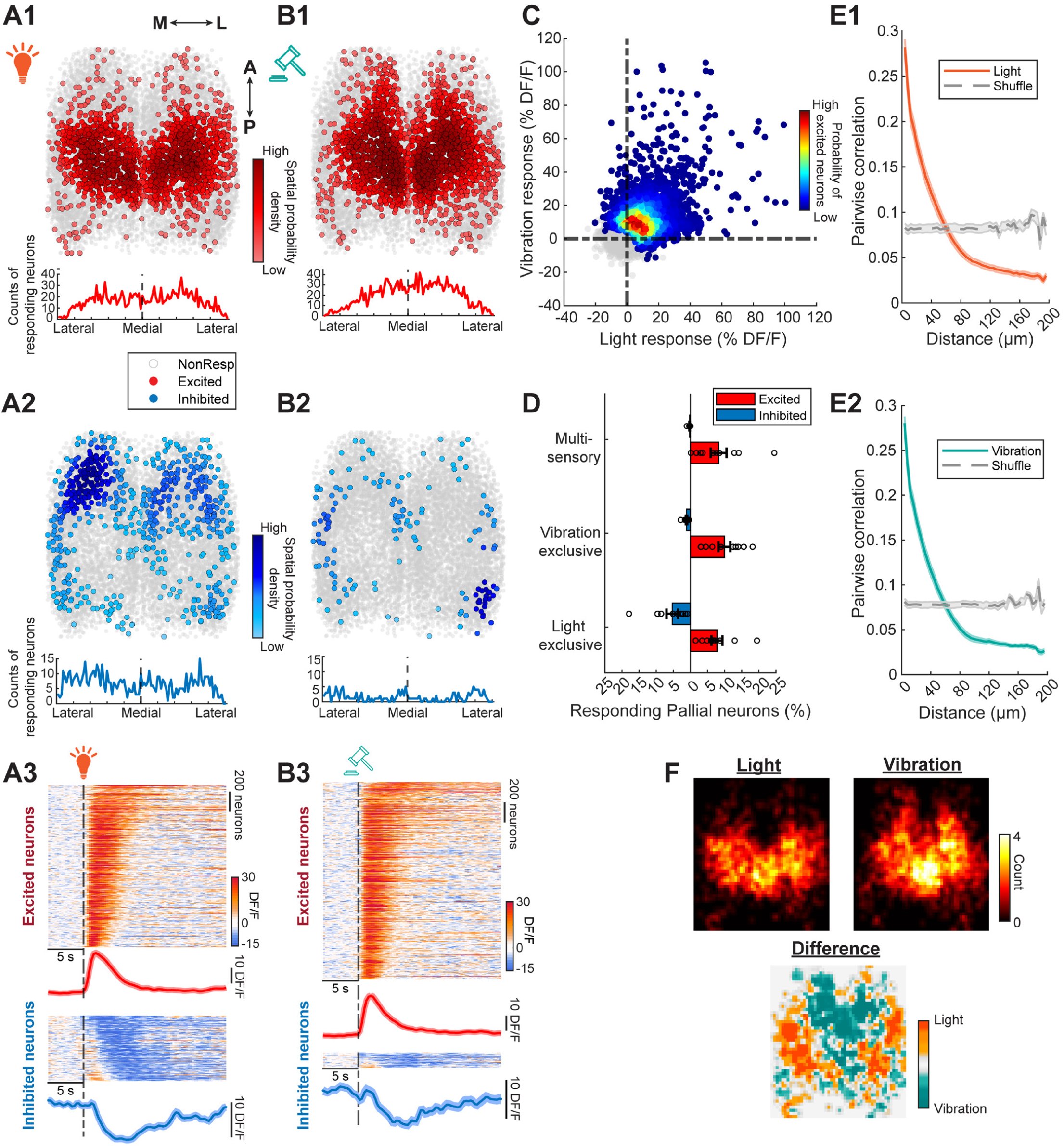
Sensory-evoked pallial responses are heterogeneous and topographically organized. **A-B**. Overlaid 2D reconstruction of the excited (in red, A1-B1) and inhibited (in blue, A2-B2) pallial neurons from all spatially aligned fish, in response to light (A) and vibration (B) stimulation, in vivo. The color gradient represents the spatial probability density of the responding pallial neurons. Non-responding pallial neurons are in gray. Time courses of the pallial neurons’ calcium signals in response to light (A3) and vibrations (B3). Warm colors indicate increased neural activity; cold colors indicate decreased activity. Mean calcium signals are shown at the bottom of each heatmap; shades indicate the standard error of the mean (N = 10 fish). **C**. Mean responses of all individual pallial neurons upon light and vibration stimulation. The probability density of the pallial neurons with significant excitation upon light or vibration stimulation is color-coded. Non-excited neurons are shown as gray dots. **D**. The fraction of responding pallial neurons excited (red) and inhibited (blue) by either light or vibration (exclusive), or by both stimuli (multi-sensory), in individual fish. **E**. The pairwise correlation of the pallial neurons’ responses to light (E1) and vibration (E2) stimulation as a function of the distance between pairs. The grey dashed line represents the pairwise correlations when the distances are shuffled. The shaded region denotes the standard error of the mean. (N = 10 fish). **F**. Top row: The locations of all light (left) and vibration (right) excited pallial neurons from all fish are spatially aligned and overlayed in a 2D histogram. Warmer colors denote higher neuron counts per bin. The difference between the two 2D histograms is shown at the bottom. Warm colors highlight a spatial preference for the light excited pallial neurons while green colors denote a preference for the vibration excited pallial neurons (N = 10 fish). Error bars are standard error of the mean (N = 10 fish, p > 0.05, Wilcoxon rank-sum test).

### Multi-sensory representations are more prominent in the zebrafish pallium than in the preglomerular nucleus

Our results showed that a significant fraction of pallial neurons respond to both light and vibration stimuli and can therefore be classified as multi-sensory (Fig. 7D). This raises the possibility that some pallial neurons integrate information from parallel sensory information streams. To investigate this, we systematically compared sensory integration across different layers of the PG-to-pallium circuit. Specifically, we analyzed sensory responses from all PG neurons (Fig. 8A), PG axons in the pallium (Fig. 8B), and pallial neurons (Fig. 8C). Strikingly, we observed significantly more multi-sensory responses in pallial neurons compared to the PG neurons and pallial PG axons (Fig. 8D-G), particularly in distinct telencephalic regions, notably the Dc and Dm areas (fig. S7B-E). We also assessed how similar light and vibration stimuli are encoded at different stages of the PG-to-pallium network. Multi-neuronal sensory responses in the pallium exhibited significantly higher Pearson’s correlation and cosine similarity than in the PG (Fig. 8H-J). These findings suggest that sensory selectivity decreases from the thalamic-like PG to the cortical-like pallium.

**Fig. 8.**
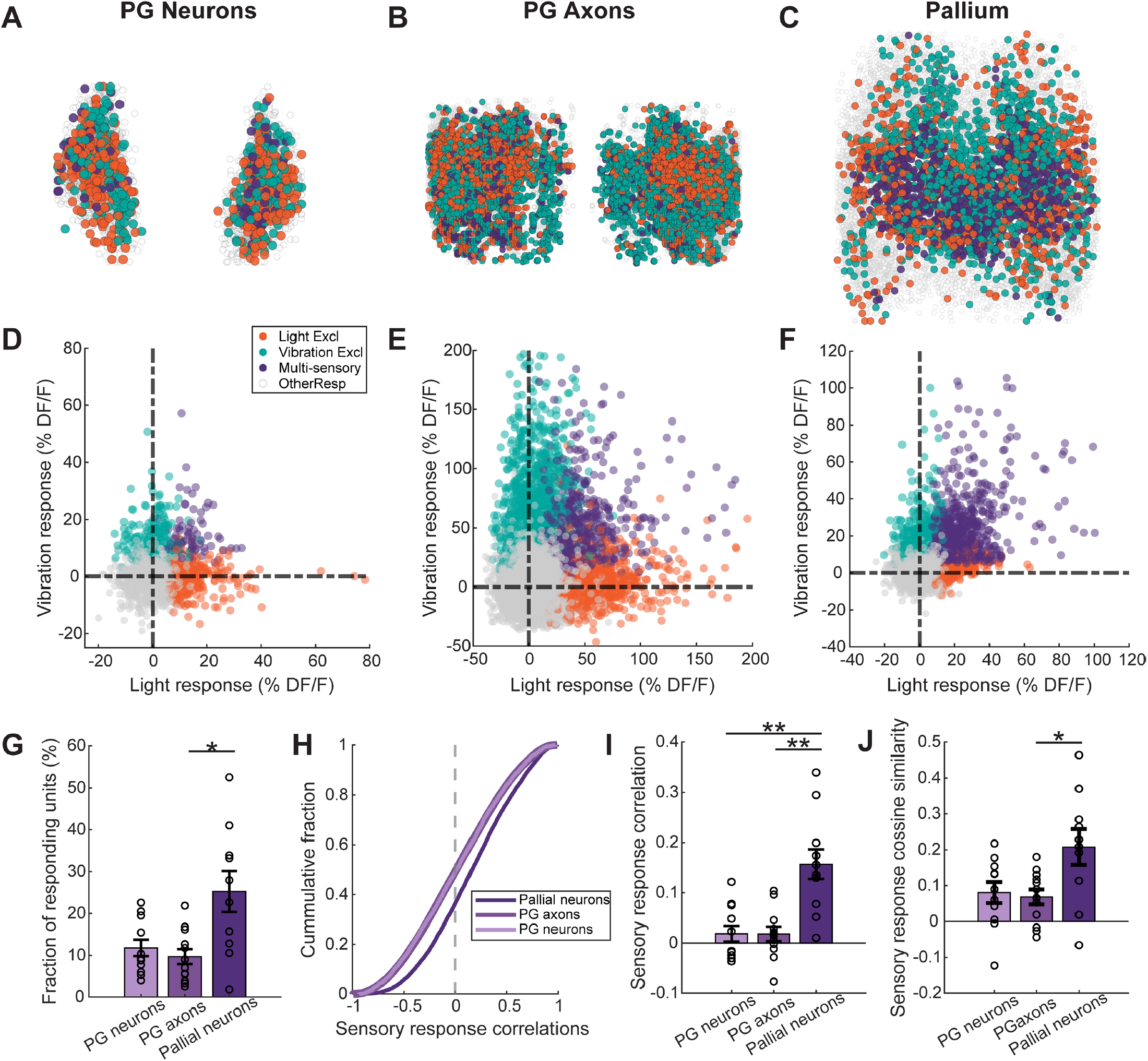
Sensory representations become less selective as they are relayed from the preglomerular nucleus to the pallium. **A-C**. 2D reconstruction of the excited PG neurons (A; N = 11 fish), PG axons (B; N = 12 fish), and pallial neurons (C; N = 10 fish) color-coded by their sensory selectivity (orange for light exclusive, teal for vibration exclusive, purple for multi-sensory responding neurons, and gray for all other neurons **D-F**. Scatter plot of the mean sensory responses of the PG neurons (D), PG axons (E), pallial neurons (F), same color-code as in A-C. **G**. The fraction of multi-sensory responding neurons/units from all datasets across the PG to pallium. **H**. Cumulative distribution of the sensory response correlations for all neurons/units across datasets (dark purple: pallial neurons, purple: PG axons, light purple: PG neurons). **I**. Mean correlations between the multi-neuronal sensory representations across the different datasets. **J**. Mean cosine similarity between the multi-neuronal sensory representations across the different datasets. Error bars denote the standard error the mean. (*p-value< 0.05, **p-value < 0.01 & ***p-value < 0.001, Wilcoxon rank-sum test).

### Hierarchical sensory processing in the zebrafish pallium is topographically organized

Our results revealed increasing complexity in how sensory information is processed from the PG to the pallium. Nevertheless, we also observed that even within the pallium, there are both simple neurons, responding exclusively to either light or vibrations, and multi-sensory neurons that responded to both stimuli. We next asked how these different categories of pallial neurons are spatially organized within the pallium. We found that neurons responding exclusively to light or vibrations are primarily located in distinct pallial zones: Dl and Dm, respectively (Fig. 9A–B), resembling the parcellation of the mammalian sensory cortices, which receive modality-specific input from distinct thalamic nuclei (2, 14). In contrast, multi-sensory pallial neurons are located in the central-medial pallium, primarily in the posterior Dc and Dm regions (Fig. 9C).

**Fig. 9.**
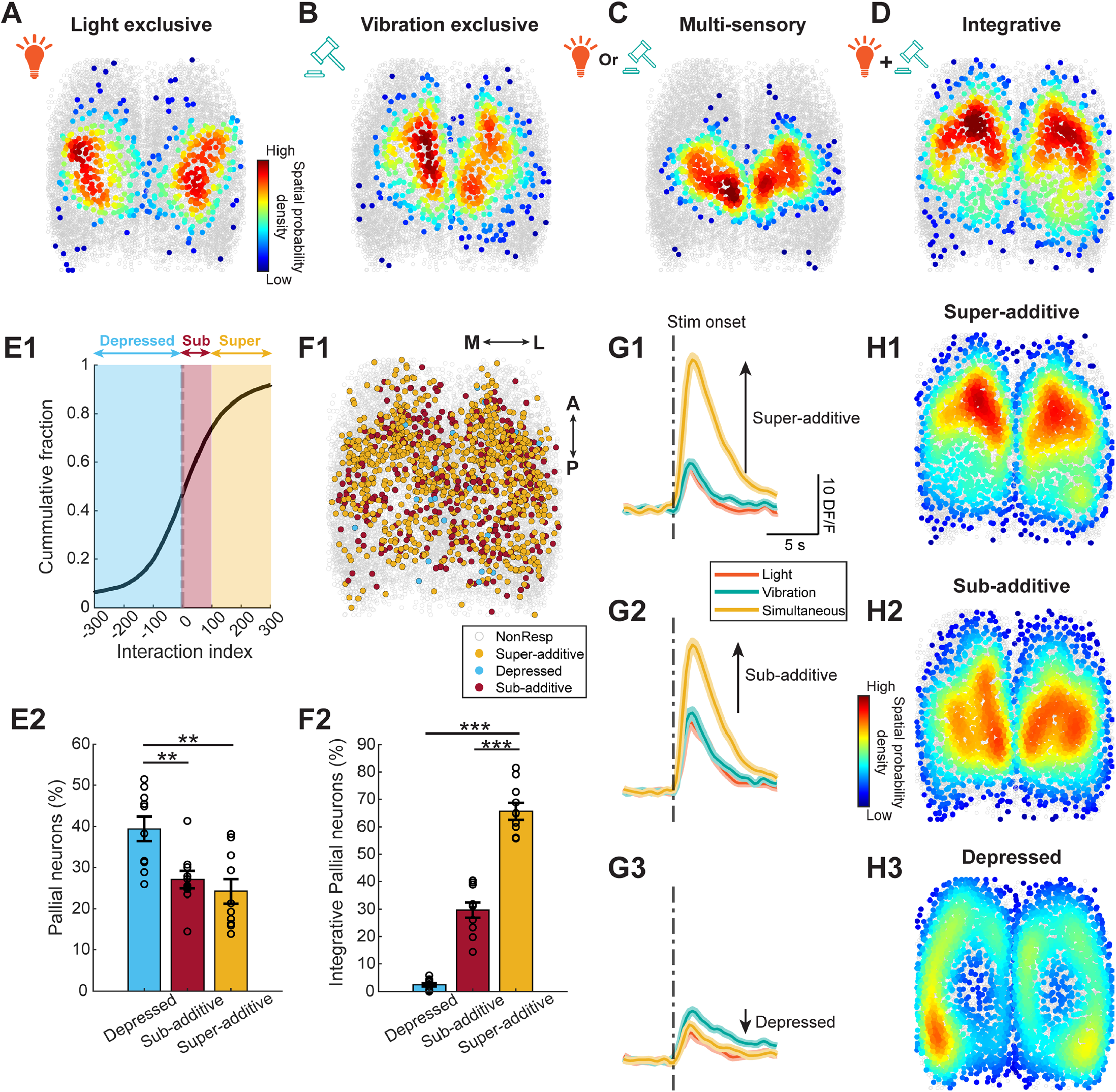
Hierarchies of sensory representations are topographically organized across the zebrafish pallium. **A-D**. Overlaid 2D reconstruction of the excited pallial neurons from all spatially aligned fish, that are exclusively responding to light (A) or vibration (B) stimulation, multi-sensory pallial neurons responding to both light and vibrations (C), and integrative pallial neurons that did not respond to individual stimuli when presented alone, but only responded when both light and vibrations are delivered simultaneously (D). The color gradient represents the spatial probability density of the excited pallial neurons. All other pallial neurons are in gray (N = 10 fish). **E1**. Cumulative distribution of the interaction index for all pallial neurons. **E2**. The fraction of pallial neurons exhibiting additivity or depression in each individual fish (N = 10 fish). **F1**. 2D reconstruction of the previously identified integrative pallial neurons classified by their interaction index, yellow for super-additivity, blue for depressed, red for sub-additivity and gray for all other neurons. **F2**. The fraction of integrative pallial neurons exhibiting additivity or depression in each individual fish. **G**. Mean responses of the pallial neurons exhibiting super-additivity (G1), sub-additivity (G2) and depression (G3) following the simultaneous delivery of both light and vibration stimuli (gold) when compared to the individual presentation of light (orange) and vibration (teal) stimuli. **H**. 2D reconstruction of the pallial neurons exhibiting super-additivity (H1), sub-additivity (H2) and depression (H3). The color gradient represents the spatial probability density of the classified pallial neurons. All other pallial neurons are in gray (N = 10 fish). Error bars denote the standard error the mean. (**p-value < 0.01 & ***p-value < 0.001, Wilcoxon rank-sum test).

After observing the increasing complexity of topographically organized sensory representations in the zebrafish pallium, we further examined the neural computation underlying sensory integration. To do this, we designed a cross-modal stimulation (105) experiment in which both light and vibration stimuli were delivered simultaneously (fig. S8A–D). This allowed us to identify pallial neurons that did not respond to either light or vibration when delivered alone but responded only to their simultaneous presentation. We henceforth referred to these as “integrative” neurons. Strikingly, integrative neurons represented the largest fraction of pallial neurons compared to other functionally defined populations (fig. S8E). Moreover, they were localized to a distinct anterior-central region of the pallium, the anterior Dc, which is spatially separate from regions containing the light-exclusive, vibration-exclusive, or multi-sensory neurons. These findings highlight a hierarchy of sensory computations in the zebrafish pallium, reflecting an increased complexity of information processing.

One could argue that these coincidence detector neurons in the pallium arise due to a thresholding or sensitivity effect. For example, when light and vibration stimuli are linearly combined, they may elicit detectable responses in less sensitive pallial neurons, which could then be mistakenly classified as nonlinear integrative neurons. To address this possibility, we adopted a simple index allowing us to quantify the nonlinerities of the multi-sensory integration (90, 106). According to this index, if a neuron’s response to coincidental stimuli exceeds the linear sum of its responses to each stimulus alone, it is classified as “super-additive” (Fig. 9E-F, G1). If the coincident response is weaker than the strongest individual response, it is classified as “depressed” (Fig.9E-F,G3). All other neurons are classified as “sub-additive” (Fig. 9E-F, G2). When we mapped the spatial distribution of these different categories of pallial neurons (Fig. 9G), a striking topographic organization emerged. Super-additive neurons were located primarily in the anterior-central pallial regions (Fig. 9H1, fig. S8F1), substantially overlapping with the spatial distribution of the integrative neurons (Fig. 9D). In fact, the majority of integrative neurons were classified as super-additive (Fig. 9F) highlighting the overlap between these populations. In contrast, sub-additive neurons were found in more posterior regions (Fig. 9H2, fig. S8F2). Depressed neurons that are the largest fraction of pallial neurons (Fig. 9E2) were located along the most anterior, posterior and lateral margins of the pallium (Fig. 9H3, fig. S8F3). Altogether, our results reveal a previously unknown functional hierarchy of sensory computations in the zebrafish pallium with strong topographical regionalization.

## DISCUSSION

### How does PG anatomy, connectivity and function relate to vertebrate thalamus?

The first order thalamus is the primary source of sensory information for the mammalian cortex (1, 14, 47). In zebrafish, the thalamus proper is known to be also involved in various sensory computations (107–112). Yet, our anatomical findings demonstrate that, although the zebrafish “thalamus proper” contains few retrogradely labeled neurons upon tracer injections into the pallium, the preglomerular complex (PG) is the predominant source of extra-telencephalic input to the pallium. This observation aligns with previous studies in teleosts (37, 40, 91, 113–116). So how does the teleost PG compare to the thalamic structures in other vertebrates? Like the mammalian thalamus (14, 95), we found that the zebrafish PG is composed primarily of glutamatergic neurons, with a smaller population of GABAergic neurons. Notably, only the glutamatergic PG neurons project to the pallium, mirroring the mammalian thalamic relay neurons (14, 95). This suggests that the GABAergic PG neurons likely function as local interneurons or thalamic reticular-like neurons (3, 56, 117, 118), mediating lateral interactions among PG neurons or modulating the local PG activity. Some support for inhibitory interactions within PG comes from anti-correlated ongoing activity between PG nuclei (e.g., Fig. 2B, clusters 1 vs. 3) and from inhibitory sensory responses observed in PG (Fig. 3). Furthermore, we showed that PG comprises of multiple functional ensembles (or clusters), reflecting the functional heterogeneity seen in the mammalian thalamus (6, 13, 46, 47, 119–121). This suggests that the zebrafish PG likely contains multiple subnuclei specialized for distinct functions, as suggested by earlier anatomical and functional work in other teleosts (41, 81, 122–124). Such functional regionalization is further supported by our observation that neighboring PG neurons exhibit more similar ongoing and sensory-driven activity patterns (Fig. 2–3). Additionally, we found that the anterolateral and posteromedial PG neurons are differentially retrogradely labeled following localized tracer injections into the Dm and Dl pallial zones, and show distinct preferences for visual versus vibrational stimuli. Dissecting the circuit architecture of the zebrafish PG will be essential for understanding how its subnuclei interact with each other and whether, like the mammalian thalamus, the PG is more than just a relay.

Unlike some first-order thalamic nuclei in mammals, the zebrafish PG receives inputs from multiple mesencephalic and diencephalic regions, consistent with findings in other teleost fish (114, 122, 125, 126). Specifically, we demonstrated that the optic tectum (visual (108, 126, 127) and the torus semicircularis (auditory and lateral line; (128–130)), the teleost homologs of the superior and inferior colliculi, are the most prominent sources of input to the PG. The dominance of these collicular inputs strongly supports the idea that, in terms of input organization, the PG is analogous to the reptilian/bird nucleus rotundus (50, 56) and the mammalian pulvinar nucleus (20, 24, 50). We found that PG projects primarily to the pallial regions Dm, Dl, and Dc, which are considered analogous to the mammalian amygdala (69, 131–133), hippocampus (66, 79, 115, 134–136), and cortex (37, 64), respectively. In birds and reptiles, the nucleus rotundus projects to the entopallium (24, 54–57, 59), while the avian “thalamus proper” targets the hyperpallium (or wulst), forming parallel ascending sensory pathways. In mammals, the pulvinar nucleus projects to several higher-order cortical areas (e.g., V2, V4, MT, parietal, temporal, frontal; (52, 137)), as well as to the amygdala (52, 138), reminiscent of the PG projections to Dc and Dm. The pulvinar also receives feedback from multiple higher-order cortical areas and is therefore classified as a higher-order thalamic structure (14, 20, 52, 139). In our study, we also observed retrogradely labeled pallial neurons following tracer injections into PG, suggesting reciprocal connectivity between PG and pallium resembling the mammalian cortico-thalamic loops (2, 95, 140). Understanding how the pallial activity modulates PG and whether these pallial–PG loops support functions such as integration across pallial regions or across sensory modalities, as seen in mammals, remains to be determined.

Finally, it appears that mammals, birds, reptiles, and fish have all evolved two parallel cortical input pathways: thalamofugal and tectofugal connections. In terms of visual processing thalamofugal inputs are more dominant in mammals (14, 50), whereas in birds and reptiles, both pathways are prominent (50, 57, 59–61). In zebrafish, the tectofugal pathway, from the tectum, via PG, to the pallium, appears to be the more dominant route. Despite the reduced thalamofugal input to the zebrafish pallium, the connectivity and computations of the PG–pallium circuits closely resemble those of mammalian thalamocortical pathways. In fact, this mimics previous findings that showed that the tectalfugal pathway in some bird species can perform similar visual processing functions as the mammalian thalamofugal pathway (50, 57, 141). It is important to emphasize that these analogies are based on the functional responses and connectivity patterns of PG described in this study. To establish robust homologies between zebrafish PG and thalamic structures in other vertebrates, a comprehensive molecular comparison of thalamic cell types is needed (26, 30, 119, 142, 143), which should be further complemented by developmental studies (13, 91, 121, 144–146).

### Hierarchies of sensory computations along the zebrafish pallium

In the amniote pallium, sensory information is topographically organized, with distinct regions dedicated to processing visual, auditory, and somatosensory information (1, 6, 11, 13, 22, 54, 147). Even in the ancestral pallium of lampreys, sensory specific topography has been observed through single-unit recordings (148). Past hodological studies in teleost fish have suggested that the lateral pallium (Dl) primarily encodes visual information, while the medial pallium (Dm) processes auditory and gustatory inputs (41, 122, 149). Here, we demonstrated that light and vibrational stimuli evoke distinct responses in the zebrafish Dl and Dm, respectively. Our PG axonal imaging data (Fig. 5) further revealed that these selective pallial responses were driven by specific PG axonal innervation patterns targeting these pallial regions. Such modality-specific pallial innervation by the PG is comparable to the sensory-selective thalamic projections to the mammalian cortex and their corresponding cortical recipient cells (1, 13, 17, 46, 150).

Yet, the zebrafish pallium does not simply mirror the sensory information it receives from the PG, but it transforms this input across distinct computational layers that are topographically organized. Firstly, our results reveal a striking decrease in sensory selectivity as information is transmitted from the PG axonal terminals to the pallial neurons (Fig. 7–8). In fact, we identified a specific central pallial region, the posterior Dc (Fig. 9C; Supplementary Fig. 7E), as the primary zone containing multi-sensory neurons with mixed selectivity. This organization closely resembles the presence of multi-sensory, mixed-selectivitive neurons observed across the mammalian cortex (8, 15, 17, 105, 151–158), and highlight multi-sensory integration as an important feature of the vertebrate pallium.

Beyond this decrease in sensory selectivity from the PG to the pallium, we also uncovered a remarkable degree of topographically organized nonlinearities in pallial sensory processing. For example, a substantial population of neurons in the anterior-central pallium (anterior-Dc) function as coincidence detectors, responding only when both light and vibration stimuli were presented simultaneously (Fig. 9D). These same neurons also exhibited nonlinear summation and amplification of responses during simultaneous light and vibration stimulation (Fig. 9G1, 9H1). We also found that most neurons in the anterior and lateral pallium show depressed responses, when both light and vibration were delivered together, suggesting stimulus competition. Altogether, these findings highlight the zebrafish pallium as a nonlinear transformer of incoming sensory information. Such nonlinear sensory computations may help explain recent findings of neurons involved in learning (69, 73, 74, 159) and spatial navigation (77–79, 135, 160) located across several pallial regions, including Dc. Our findings are also in line with a plethora of literature on learning related nonlinearities across the mammalian cortex (15, 16, 161–164).

In mammals, these nonlinearities can be generated by distinct circuit motifs (1, 2, 6, 16, 17, 165–168) but where and how do these pallial nonlinearities emerge in teleosts? Previous anatomical studies of the teleost pallium have highlighted a rich diversity of architectural features, including recurrent (94, 134), feedforward/feedback (40, 41, 115, 122, 149, 169, 170), and interhemispheric (40, 93, 169) connections. These connectivity motifs are also well-documented in other amniotes including the bird (22–24) and reptilian pallium (27, 29, 171) suggesting that these architectural motifs are highly conserved across vertebrates. Additionally, we observed bilateral pallial responses following ipsilateral PG stimulation, indicating the presence of excitatory cross-hemispheric connections (Fig. 6). In fact, PG stimulation not only activated neurons near the PG axon terminals in the pallium but also triggered activity that spread to the anterior pallium, including areas lacking direct PG innervations. This multi-synaptic propagation of PG inputs is further supported by our observation of reduced reliability in PG-evoked responses across contralateral and anterior pallial regions (Fig. 6E–F). Moreover, ~40% of pallial neurons exhibited competitive suppression when light and vibration stimuli were presented simultaneously (Fig. 8E). Given the predominant glutamatergic nature of the zebrafish pallium (36, 74, 172), this stimulus competition is potentially mediated by feedback connections between glutamatergic pallial neurons and GABAergic subpallial neurons (63, 170, 173, 174). Consistent with this, stimulation of ipsilateral glutamatergic PG-to-pallium projections not only evoked widespread excitation but also induced prominent inhibition in both hemispheres, likely via subpallial GABAergic circuits. While this study focused on the function and anatomy of PG-to-pallium connections, our findings underscore the need for a comprehensive mapping of the anatomical, synaptic, and functional connectivity within the zebrafish pallium. With its small, optically accessible brain and powerful genetic toolkit, the zebrafish is well-positioned to become one of the first vertebrates to achieve a complete pallial connectome, through a combination of electron microscopy (175, 176) and optogenetic approaches (177, 178), following the major advances made in the fruit fly connectome (179, 180).

### Functional topography and regionalization of the vertebrate pallium

Sensory-motor computations in the zebrafish midbrain and hindbrain have been extensively studied (177, 181–189). However, how (non-olfactory) sensory information is received, represented, and processed in the zebrafish pallium, the ancestral homolog of the mammalian cortex, remains less understood. In this study, we demonstrated that despite the near absence of thalamofugal inputs, the zebrafish pallium has evolved convergent architectures and computational hierarchies using tectofugal inputs relayed through the PG, a hub-like nucleus analogous to the reptilian and avian nucleus rotundus and the mammalian pulvinar nucleus. Our findings reveal that specific sensory computations in the zebrafish pallium (i.e., modality-specific responses, multi-sensory integration, and coincidence detection) are spatially segregated and topographically organized (Fig. 10). We also showed that this pallial topography exhibits a striking hierarchy, with increasingly complex and nonlinear response patterns emerging along the posterior-anterior axis. On one hand, this regionalization resembles the nuclear, non-laminated structure of the avian pallium (23, 24, 58). On the other hand, the gradient of increasing nonlinearity from posterior to anterior pallial regions is reminiscent of the functional hierarchy observed in the mammalian cortex, where simpler computations occur in the posterior sensory areas while more complex, and integrative processing take place in the anterior higher-order cortices of mammals (1, 7, 8, 15, 190) and other amniotes (27, 147). Together, our results and previous findings highlight regionalization as a conserved and fundamental feature of the vertebrate pallium. This regionalization likely supports the pallium’s role as a parallel processor capable of performing multiple operations simultaneously. Whether this functional regionalization of the zebrafish pallium is supported by distinct molecular expression patterns is still unknown. Recent spatial transcriptomic studies have mapped the molecular topography of pallium in various vertebrates (25, 26, 28, 30, 143), including mammals (142, 191) pallium with unprecedented resolution. We anticipate that generating a comparable molecular atlas of the zebrafish pallium will enable novel cross-species comparisons of the molecular, architectural, and functional features of the different pallial neurons across the vertebrate taxa.

**Fig. 10.**
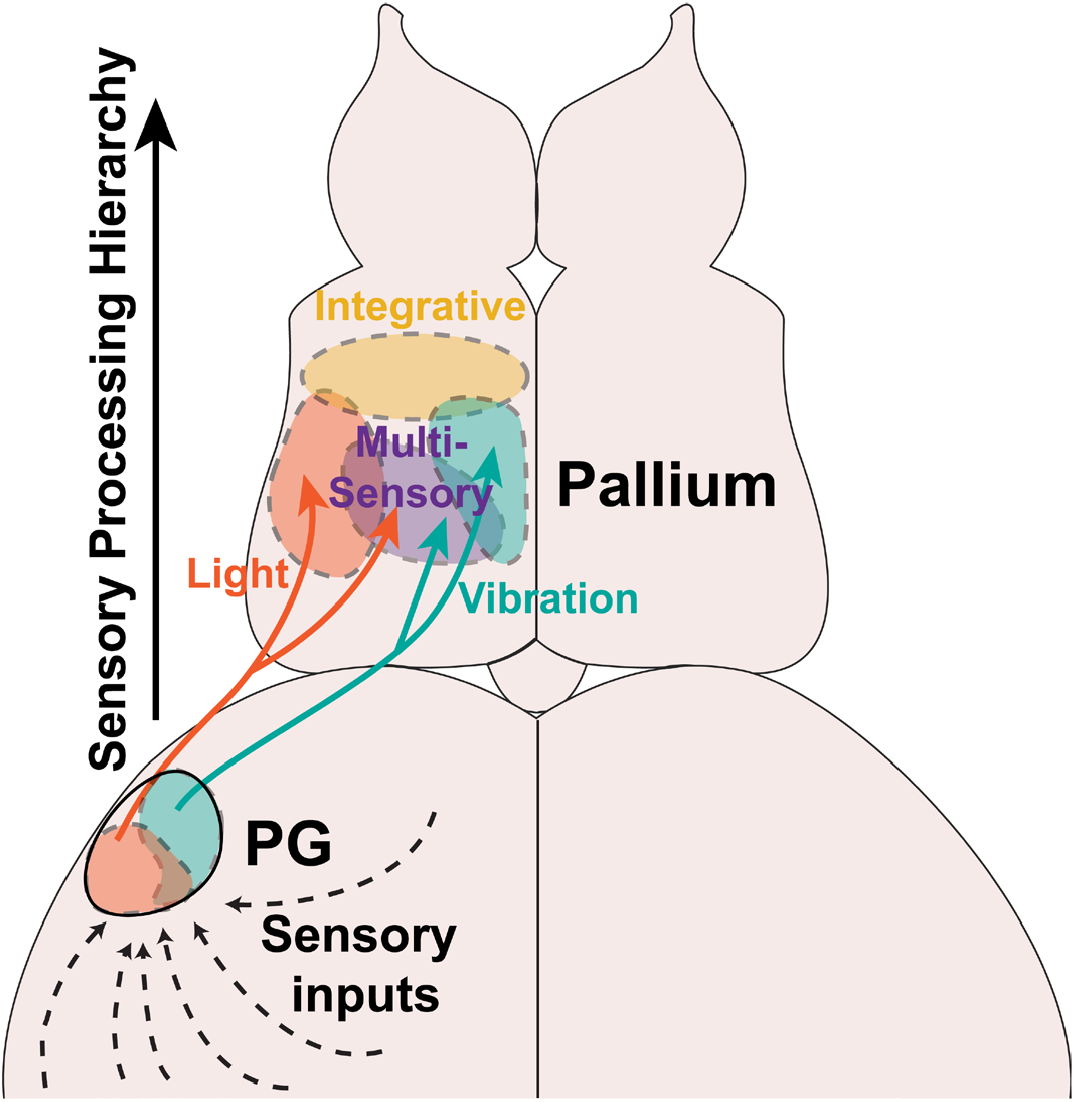
Working model of sensory transformations from the preglomerular nucleus to the zebrafish pallium. The preglomerular nucleus (PG) is the primary sensory input channel, delivering visual and vibrational information to the zebrafish pallium. Representations of these stimuli are selectively encoded by distinct populations both within the PG and in its axonal projections to the pallium. The pallial PG projections are topographically organized into distinct zones that preferentially respond to specific sensory modalities. In the pallium, sensory responses are less selective than those in the PG, exhibit diverse heterogeneity, and are hierarchically organized with notable topographic structure. Pallial neurons with selective sensory responses are arranged into distinct zones along the medial-lateral axis. Multi-sensory pallial neurons are prominent in the transitional regions between these zones. Integrative pallial neurons, which exhibit nonlinear (super-additive) responses and respond only when both visual and vibrational stimuli are delivered simultaneously, are localized to a distinct anterior pallial region. Hence, the complexity of sensory representations increases from the posterior to the anterior zebrafish pallium in a hierarchical manner.

## ACKNOWLEDGEMENTS

We thank M. Ahrens (HHMI, Janelia Farm, USA), S. Higashijima (Okazaki Institute for Integrative Bioscience, Japan) for the transgenic fish lines; We also thank S. Eggen, F. Acuña-Hinrichsen, V. Nguyen, and our fish facility support team for technical assistance. We thank Leonard Maler (University of Ottawa, Canada), and the Yaksi lab for stimulating discussions. This work was funded by the JSPS KAKENHI grant (JP24K02008) to KK, the European Horizon Marie-Curie individual postdoctoral fellowship (grant ID: 101066743) to A-TT, the NFR FRIPRO research grants 239973 and 314212 to EY and RCN Centres of Excellence scheme, project number 332640. Work in the EY laboratory is funded by the Kavli Institute for Systems Neuroscience at the Norwegian University of Science and Technology.

## AUTHOR CONTRIBUTIONS

Conceptualization: A-TT, EY; Methodology: A-TT, AO, IDCB, SK, FC, BS, KK, EY; Investigation: A-TT, AO, IDCB, SK, FC, BS, EY; Visualization: A-TT, AO, IDCB, EY; Funding acquisition: A-TT, KK, EY; Project administration: EY; Supervision: A-TT, EY; Writing – original draft: A-TT, EY; Writing – review & editing: all authors.

## INCLUSION AND ETHICS STATEMENT

Our team includes researchers from diverse origins and backgrounds. All experimental procedures performed on zebrafish larvae and juveniles were in accordance with the Directive 2010/63/EU of the European Parliament and the Council of the European Union and approved by the Norwegian Food Safety Authorities. Animals from both sexes were used in this study.

## DATA AND CODE AVAILABILITY

Processed calcium imaging data will be deposited to an open-access data repository before publication. The analysis codes and figure-making associated codes are available online on Github at: https://github.com/yaksilab/PG-Pallium_paper

## MATERIALS AND METHODS

### Materials availability

This study did not generate new unique reagents.

### Zebrafish husbandry and strains

Juvenile (21 – 28 days post fertilization) were used for the majority of the experiments except otherwise stated. All zebrafish were kept in 3.5 Ltanks and maintained at a temperature of 28°C, pH 7.2, ~700 μSiemens. Fish were kept on a 14:10 day/night cycle and all fish maintenance procedures have been approved by the Norwegian food and safety authority (NFSA). Zebrafish at this age are not sexually matured and hence, do have a gender yet. Fish from the following transgenic lies were used for the experiments: the *Tg(vglut2a:DsRed;gad1b:GFP)* which was generated by crossing the *Tg(vglut2a:DSred)* (96); *Tg(gad1b:GFP)* (97), the *Tg(elavl3:GCaMP6s)* (100), *Tg(elavl3:H2B-GCaMP6s)* (100), *Tg(gSAIzGFFD707A: Gal4;UAS:GCaMP6s)* which was created by crossing fish from the *Tg(gSAIzGFFD707A:Gal4)* and *Tg(USA:GCaMP6s)* (192) lines. The *Tg(gSAIzGFFD707A:Gal4)* was generated at the National Institute of Genetics (Mishima, Japan) using the Tol2 transposon-based gene trap and enhancer trap constructs (69, 193, 194).

### Explant preparations

The explants were prepared as previously described in (70). In short, the fish were first anesthetized in ice-cooled artificial fish water (AFW, 1.2g of marine salt in 20L of distilled water) and then euthanized by decapitation in oxygenated (Carbogen, 95% O2/ 5% CO2, Linde Corporation) artificial cerebrospinal fluid (ACSF). The ACSF was prepared by dissolving the following salts in distilled (reversed-osmosis, RO) water: 131 mM NaCl, 2 mM KCl, 1.23 mM KH2PO4, 2 mM MgSO47H2O, 10 mM glucose, 2.5 mM CaCl2, and 20 mM NaHCO3 (93, 195). After euthanization, the underlying muscle tissue, jaws and eyes were first removed. Afterwards, the skin, dura and bones were carefully removed to provide the intact dissected brain with free access to the oxygenated ACSF. The extracted brain was then mounted to a small petri dish coated with Sylgard (World Precision Instruments) using tungsten pins prior to being transferred to the imaging setup (either epifluorescent or two-photon setups, see below) where the brain was constantly perfused with oxygenated ACSF. They were then used for either neurotracer or calcium imaging experiments (*ex vivo* Ca^2+^ experiments section below).

### Neurotracer injections

Anatomical tracing experiments were performed in juvenile zebrafish explants of the *Tg(elavl3:GCaMP6s)* and *Tg(gad1b:GFP)* fishlines. For these experiments, brains were visualized using a brightfield/epifluorescent microscope (Olympus BX51WL) controlled by a manual stage. Fluorescence from the dye was visualized using a green LED (530 nm) (M530L3, ThorLabs). Both a 5x air objective (Olympus, MPlanFLN NA 0.15) and a 10x water immersion objective (Olympus, MPlanFLN NA 0.3) were used to visualize the brain explants. Tetramethylrhodamine dextran dyes (3000 MW, Thermofischer) were used to label cell bodies and projections of interests. Given that the dye is charged, we used iontophoresis to perform localized injection of the dye in our structure of interests. The neurotracer dye was loaded into borosilicate glass pipettes (1.00 mm; World Precision Instruments) which were pulled using a horizontal puller (P-2000, Shutter Instruments). Using a silver-wired electrode, we applied small electrical pulses at various intervals (Master8, A.M.P.I.) and amplitudes using a stimulating unit (ISO-Flex, A.M.P.I.) to control the size of the injection under the epifluorescence microscope. Small injections were performed using 200-400 pulses of 25 ms at an interval of 1Hz using currents of 0.2 mA. In contrast, big injections were done using 600-800 pulses with the same frequency and amplitude. Following the neurotracer injections, the explants were kept perfused with oxygenated ACSF for 3-4 hours before being incubated in the same ACSF at 4°C overnight. The samples were then fixed in 4% paraformaldehyde for 4-6 hours. Following the tissue fixation, the juvenile zebrafish brain explants underwent a tissue clearing process using the recommended protocol from the Binaree Tissue Clearing Kit (#HRTC-012, Binaree). In short, the tissue was incubated in the Binaree Starting solution at 4°C overnight. On the next day, the samples were transferred to a well plate containing 500 μL of Tissue Clearing Solution A and incubated at 37°C for 24 hours. Then, the samples were washed four times with RO water while shaking at 30 rpm at 4°C for 20 minutes each. The samples were then transferred to another well plate containing 500 μL of Tissue Clearing Solution B and incubated at 37°C for another 24 hours. Finally, the tissue was washed four times again with distilled (RO) water as done previously before being mounted on a microscope slide with the provided mounting solution from Binaree.

### Confocal imaging and anatomical analysis

Once cleared, the explants were then imaged using a confocal microscope (Examiner Z1 LSM 880 confocal microscope, Zeiss) and using either a 10x (Zeiss, Plan-Apochromat NA 0.45) or 20x objective (Zeiss, Plan-Apochromat, NA 0.8). Z-stack images were acquired using Zen 3.9 (Zeiss) and then further analyzed using either ImageJ/Fiji or Imaris 4.0 (Oxford Instruments). Cell counts were manually done in Imaris 4.0. We based our anatomical analysis on the anatomical landmarks identified in (40, 70) and the adult zebrafish brain atlas (AZBA) (62).

### *In Vivo* two-photon Ca^2+^ imaging

For *vivo* experiments, a two-photon microscope system (Scientifica) with a 20x water immersion objective (Zeiss 7 MP, W Plan-Apochromat NA 1.0) was used. Excitation was done using a mode-locked Ti:Saphire laser (MaiTai, Spectra Physics) tuned to 920 nm. Volumetric in vivo recordings were done across 8 planes using a Piezo (Physik Instrumente) at an acquisition rate of 2.2 – 3.4Hz per volume. Image sizes of 1536 x 850 pixels were used for the PG recordings while image sizes of 1536 x 600 pixels were used for the telencephalic recordings. All recordings were done using the SciScan software package (LabView).

For *in vivo* two-photon calcium imaging experiments, the juvenile zebrafish were first embedded in agarose as per (70, 71, 76, 90). In brief, the fish were embedded in low-melting point (LMP, Fisher Scientific) agarose (2.5% in AFW) in a recording dish (Fluorodish, World Precision Instruments) and waited for 20 minutes for the agarose to solidify. Afterwards, a triangular section of agarose around the nose, mouth and another triangular section encompassing the tail were removed with a scalpel. The embedded fish was then transferred to the recording setup where the recording dish was constantly perfused with heated (28 °C, Warner Instrument Corporation), oxygenated (Carbogen) AFW for 30 minutes prior to the recording.

Prior to sensory stimulation, ongoing spontaneous activity was recorded for 10 minutes. Afterwards, sensory stimulation (either a red-light flash or a mechanical vibration) was presented to the fish at different intervals, using parameters from previous studies (71, 76, 90). The red light was produced by a red LED (LZ1-00R105, LedEngin; 630-nm wavelength), which was placed in front of the fish. The light flashes were 200 ms long and had an intensity of 0.318 mW at 635nm, at the location of the fish during these experiments. Additionally, the mechanical vibrations were produced by applying 50 ms-long 6 V current to a solenoid tapper (SparkFunElectronics, ROB-10391) generating a vibration frequency spanning between 300-10000Hz (90). Both of which were controlled by an Arduino connected to the computer and initiated using Matlab (Mathworks). These sensory stimuli were delivered either individually (Figs. 3, 5, 7, S4, S5, S7) or simultaneously (where the onsets were synchronized, Figs. 9, S8), and were all used in previous studies (71, 76, 90). Light, vibration or co-stimulation of light and vibration were repeated eight times with a interstimulus interval of 60 s. Finally, the time between stimulus conditions was 4 mins.

### *Ex vivo* Ca^2+^ imaging

For explant calcium imaging experiments, a similar two-photon microscope setup (Scientifica) conneceted to a Ti:Saphire laser (MaiTai, Spectra Physics) tuned to 920 nm was used. The same acquisition software (SciScan, LabView) was also used except that a 16x water immersion objective (Nikon, NA 0.8, LWD 3.0) was used instead. Volumetric imaging of the entire telencephalon was achieved by imaging 8 planes with an image size of 1536 x 750 pixels at a rate of 2.2-2.5Hz.

Additionally, control micro-stimulation experiments were done using an epifluorescent microscope setup similar to the neurotracer experiments. In this case, a blue (470 nm) LED (M470L3, ThorLabs) was used to excite and visualize the GCaMP fluorescence. Fluorescent images were acquired using a QImaging camera (Teledyne Photometrics) and using the Occular image acquisition software (Teledyne Photometrics). Single-plane images of the explants were acquired at 40 Hz using a camera exposure setting of 4 ms.

#### Micro-electrode stimulation

Glass pipette micro-electrodes were prepared as in (70) by first pulling borosilicate glass capillaries (1.00 mm; World Precision Instruments) using a horizontal puller (Model P-2000, Sutter Instruments). The glass electrodes had a resistance of ~10-12 MOhm. Custom bipolar electrodes were created by gluing two glass electrodes using a two-component epoxy glue (Locite) so that the glass pipette tips were within 1-2 mm of each other. After, the glass pipettes were filled with ACSF and positioned in the brain using micro-manipulators (Scientifica). PG neurons were stimulated using a train of ten short (2 ms) current pulses (30-50 μA) with an inter-stimulus interval of 60 s using a current isolator unit, a model DS3 (Digitimer). For the control experiments (fig. S6), a train of 40-50 μA current pulses were used to stimulate the neurons in the PG and adjacent brain area (~ 20 μm outside of the PG).

### Data Analysis and Quantification

Two-photon microscopy images were aligned using Suite2p (196) and then visually inspected for any aberrant motion artifacts, such as drift in the z-dimension. Only those experiments with corrected motion and drift were used for further analysis. Regions of interest (ROI) corresponding to neurons were detected using a custom semi-automatic template-matching algorithm (70, 71, 90, 101, 103). The spatial positions of the ROIs were then extracted using custom Matlab scripts and the relative change in fluorescence (ΔF/F) was calculated for each neuron. For sensory and electrical stimulation experiments, the ΔF/F was calculated based on the 5s baseline prior to each stimulation trial. In contrast, during ongoing spontaneous activity periods, the ΔF/F was calculated using a sliding baseline window of 6 mins. Afterwards, the ΔF/F signals were filtered using a low-pass filter as described in (197). The aforementioned Ca^2+^ signal processing was all done in Matlab (Mathworks) using custom scripts. Delineation of the brain regions was done manually based on anatomical landmarks identified in (40, 70) and AZBA (62). For telencephalic recordings, only the dorsal telencephalon were analyzed and considered as the zebrafish pallium. The telencephalic PG axonal detection was done using the methods described in (76, 198). In brief, raw fluorescent images were first binned before a threshold-based algorithm was used to detect axonal pixels in a manually drawn region of the binned fluorescent image. Binned pixels with relative fluorescence intensity values greater than 25% of the maximum pixel value were considered as axonal pixels. Afterwards an independent comonent analysis was made to the fluorescent images to identify axonal ROIs (76, 198) which were then manually inspected prior to further analysis.

Spontaneous ongoing neuronal activity of individual neurons was clustered into functional ensembles using the k-means clustering function in Matlab (70, 90, 101). The optimal number of clusters used for the clustering was identified using the elbow method (70, 76, 101). In brief, the elbow method calculates the sum of intra-cluster distances for each cluster element, normalized by the sum of average inter-cluster distances for the actual data (black), and simulated data (gray,100 iterations) with the same variance as the actual data but without cluster structure. This calculation was iterated for up to 30 k-means clusters. The relevant number of k-means clusters corresponds to the elbow point, where the black curve exhibits a prominent bend. This analysis was done for both PG neuronal data (fig. S4, N = 11 fish) and pallial PG axonal data (fig. S5, N = 12 fish).

To assess whether the cluster identity of these neurons was stable across time, we used the cluster fidelity index (70, 101). In short, this index measures the probability of each neuron remaining in the same k-means cluster across different time periods. These cluster fidelity values were then compared to a shuffled distribution where the cluster identities of each neuron were randomized.

To assess the functional topography of the PG and pallial recording units (neurons and axons) following sensory stimulations in vivo and the electrical micro-stimulations *ex vivo*, only responding units were considered for further analysis. These responding units were quantified as described in previous work (70, 90). In brief, neurons/axons were classified as positively responding (“excited”) when their mean responses across trials during a 5 s time window following the stimulus onset (response period) was greater than the mean plus 2.5 times the standard deviation of the baseline activity (5 s before the stimulus onset). In contrast, negatively responding (“inhibited”) neurons/axons were identified when the mean activity during the response period was smaller than the mean plus 1.5 times the standard deviation of the baseline activity.

To visualize the spatial distribution of responding units across fish, we have normalized the spatial positions along the X and Y dimensions of all identified ROIs (neurons or axonal pixels) per fish to a value between 0 and 1, that correspond to X and Y boundaries of the aligned brains. After spatial alignment, neurons from all recorded fish were then overlayed on top of each other and then visualized using a density plot based on a kernel smoothing function or as a 2D histogram. This 2D reconstruction of the responding neuron’s spatial positions was done individually for each hemisphere of the brain except for Fig. 6 where we examined the ipsi vs contralateral responses in the pallium. To quantify the topographic selectivity of our datasets, the counts from the vibration 2D histogram were then subtracted from the counts of the light 2D histogram before being smooth with a gaussian filter to create the “difference” 2D histogram.

To quantify whether the excited neurons/axons showed prominent spatial localization, we used a focality index (71). In short, the focality index ranges from 0 to 1 (0 indicating a highly random distribution and 1 indicating a highly focal distribution of). It was calculated as 1 minus the mean Euclidean distance of all excited neuronal pairs divided by the mean Euclidean distances for all neurons within one hemisphere. The average focality index of each hemisphere was then used to calculate the mean focality index per fish.

The time course traces of all responding or axonal pixels were sorted using Rastermap (199) and then visualized as heatmaps using Matlab (Mathworks).

To quantify the sensory selectivity of the responding units, we compare the mean responses vectors, which are the average response (ΔF/F) amplitudes of the excited during the response period. To examine how sensory selectivity is represented across the sensory processing hierarchy, we correlated the mean sensory response vector following light and vibration stimulation across all recorded units (Fig. 8). Units that only responded to either light or vibration were classified as exclusive neurons/axons. The units that responded to both sensory stimuli were classified as multi-sensory neurons/axons. Neurons responding only when both stimuli were presented simultaneously, but not when presented individually were classified as integrative neurons.

To futher quantify the response properties of pallial neurons (Fig. 9), we calculated an interactive index (90, 106). In brief, we measured the neuron’s average response (R) to light, vibration or co-stimulation (light and vibration) during the responding period (5 s following the stimulus onset). Next, the interactive index was calculated using the following adapted equation:

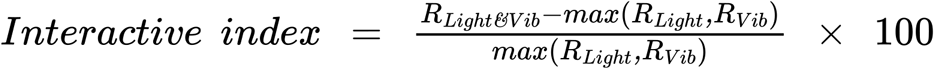

If the interactive index is >100, then the neuron was classified as being “supper-additive”, where its response to the co-stimulation was twice greater than its maximal response during individual stimulation. In contrast, if the interactive index is <0, then the neuron was classified as being “depressed” since its co-stimulation response was smaller than the maximal response during individual stimulation. An interactive index value between 0 and 100 was classified as being “sub-additive”.

All analysis figures were generated in Matlab (Mathworks) and then assembled in Illustrator (Adobe).

### Statistics

All statistical analyses were done in Matlab. Paired tests were performed using Wilcoxon signed rank test while non-paired tests were performed using the Wilcoxon rank sum test. For comparison across delineated brain regions, a Kruskal-Wallis’s test was first performed followed by posthoc Dunn’s test. All error bars represent the standard error of the mean unless specified elsewhere.

## SUPPLEMENTAL FIGURES

**fig. S1.**
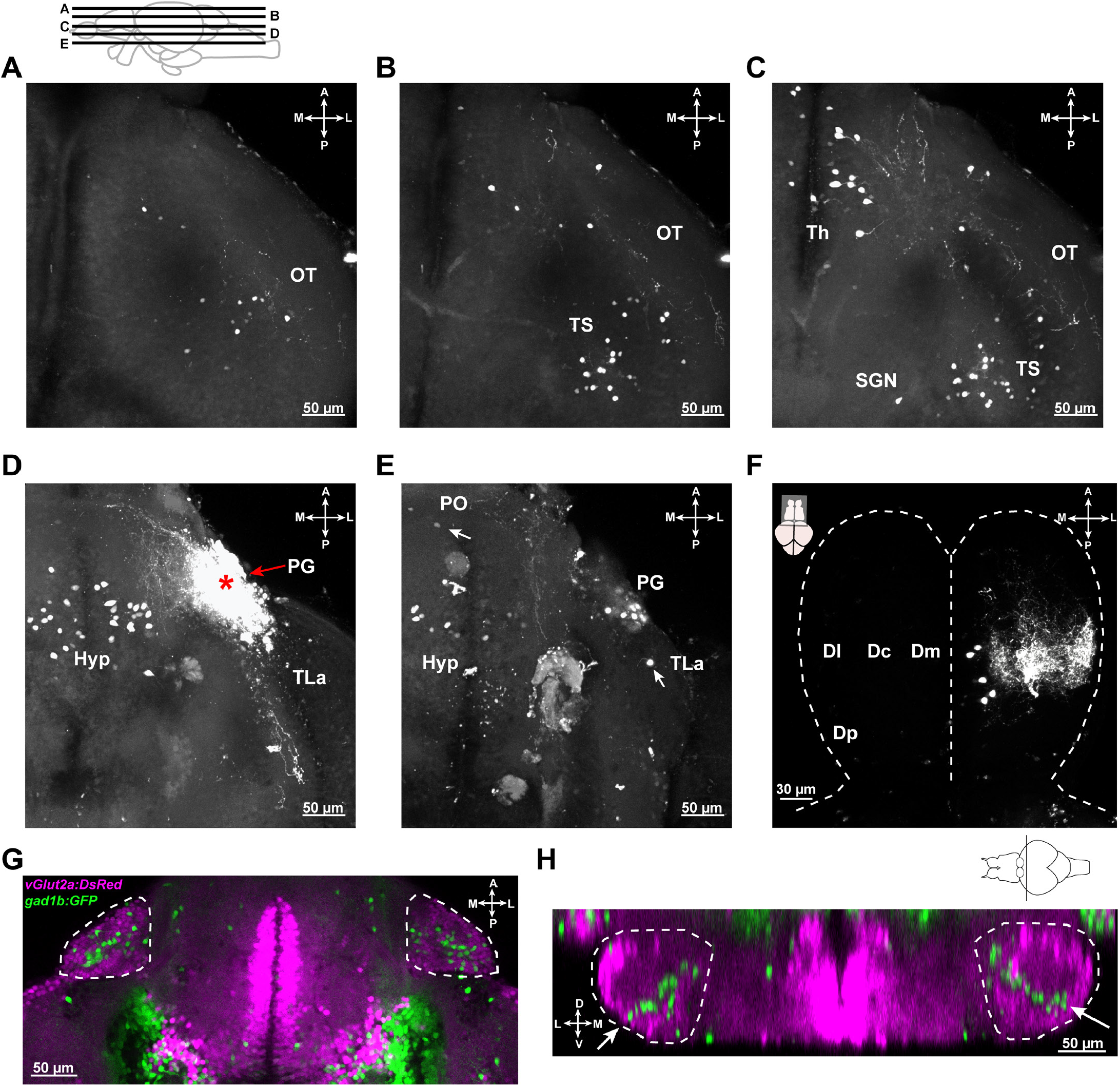
Characterization of retrogradely labeled inputs of the preglomerular nucleus and the neurotransmitter profiles of preglomerular nucleus neurons. **A-E**. Confocal microscopy images from tissue-cleared juvenile zebrafish explant containing both the diencephalon and mesencephalon illustrating examples of retrogradely labelled neurons upon neurotracer injections (TMR-Dextran) in the PG. Retrogradely labelled neurons were visible in the optic tectum (OT) in A, the torus semicircularis (TS) in B, the thalamus proper (Th), the secondary gustatory nucleus (SGN) in C, the hypothalamus (Hyp) in D, the preoptic area (PO) and the lateral torus (TLa) in E. The injection site located in the PG was shown in D. The scheme above A indicates the horizontal locations of the imaging planes corresponding to individual panels. **F**. Dorsal view of the dorsal telencephalon (pallium) showing retrogradely labelled neurons and axonal projections following a neurotracer injection in the PG. Dashed lines mark the boundaries of the telencephalon. **G**. Confocal microscopy image of both preglomerular nuclei (white dashed lines) in a *Tg(vglut2a:DsRed; gad1b:GFP)* juvenile zebrafish brain explant, horizontal view. **H**. Same as in G, but in a coronal view. The white arrows highlight the GABAergic neurons that form a band from the medial PG to the lateral-ventral PG. The scheme above H indicates the approximate location of the coronal section. Abbreviations: A: anterior, M: medial, L: lateral, P: posterior.

**fig. S2.**
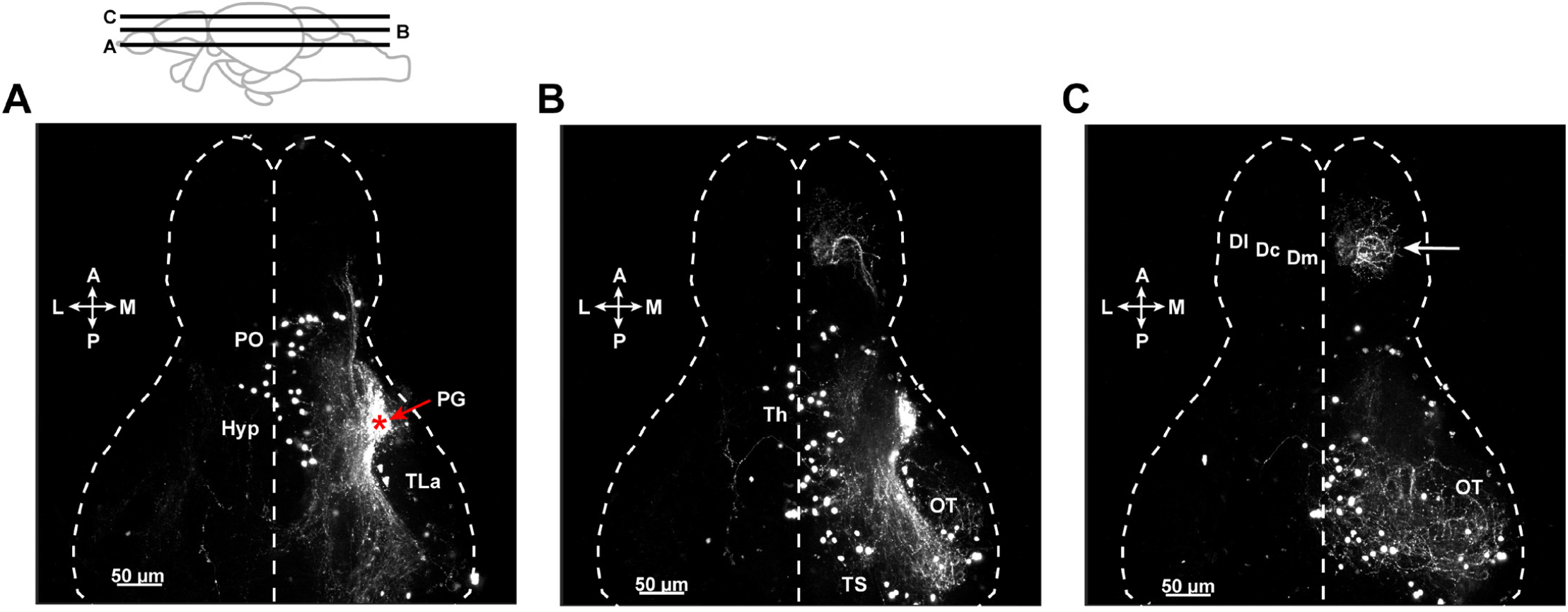
General connectivity architecture of the preglomerular nucleus outputs and inputs are present as early as in 10-14 days old zebrafish larvae. **A-C**. Confocal microscopy images from a tissue-cleared zebrafish showing the forebrain and midbrain, after a neural tracer (TMR-dextran) injection in the PG of 10-14 days old zebrafish (N = 3). The white dashed line marks the outline of the brain. **A**. Ventral imaging plane illustrating the ventral midbrain and diencephalic structures including the hypothalamus (Hyp), the preoptic areas (PO) as well as the lateral torus (TLa). The injection site, in PG, is labelled with a red star. **B**. Higher imaging plane in the same animal containing the thalamus proper (Th) and torus semicircularis (TS) and parts of the optic tectum (OT). **C**. Dorsal imaging plane in the same animal containing the dorsal telencephalic regions (Dl, Dc, Dm) and the optic tectum (OT). The arrow highlights the distribution of the PG axons projecting to the pallial regions: Dl, Dc and Dm. Scale bars are 50 μm. The illustration above A indicates the horizontal locations of the imaging planes corresponding to the individual panels.

**fig. S3.**
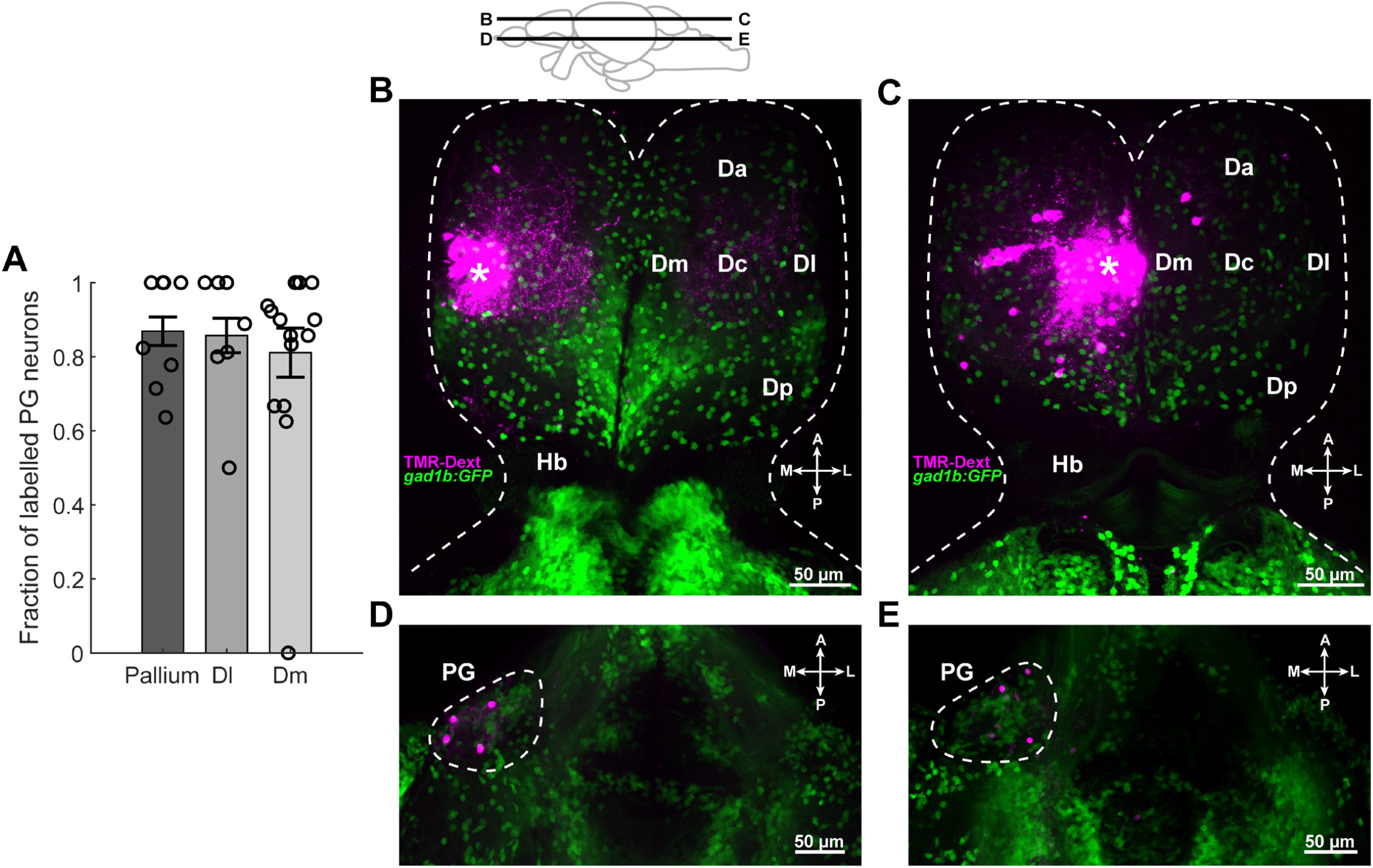
Spatially restricted neural traces injections to distinct pallial regions Dl versus Dm, retrogradely labelled distinct populations in the lateral vs medial preglomerular nucleus. **A**. The fraction of retrogradely labelled diencephalic neurons in PG versus the thalamus proper, upon broad neural tracer injections in the zebrafish pallium (N = 8 fish), specifically in Dl (N = 7 fish) and in Dm (N = 15 fish). **B-C**. Confocal microscopy images from the dorsal pallium showing examples of restricted neurotracer injections to Dl (B) and Dm (C) in Tg(gad1b:GFP) juvenile zebrafish. The white star denotes the locations of the injections. Dashed lines mark the boundaries of the telencephalon. **D-E**. Confocal images from the ventral plane illustrating the locations of retrogradely labelled PG neurons upon restricted neural tracer injections in Dl (D) and in Dm (E). Dashed lines mark the boundaries of the PG. Abbreviations: Da: dorsal anterior, Dm: dorsal medial, Dc: dorsal central, Dl: dorsal lateral, Dp: dorsal posterior, Hb: habenula, PG: preglomerular nucleus. The illustration above B indicates the locations of the horizontal imaging planes corresponding to individual panels.

**fig. S4.**
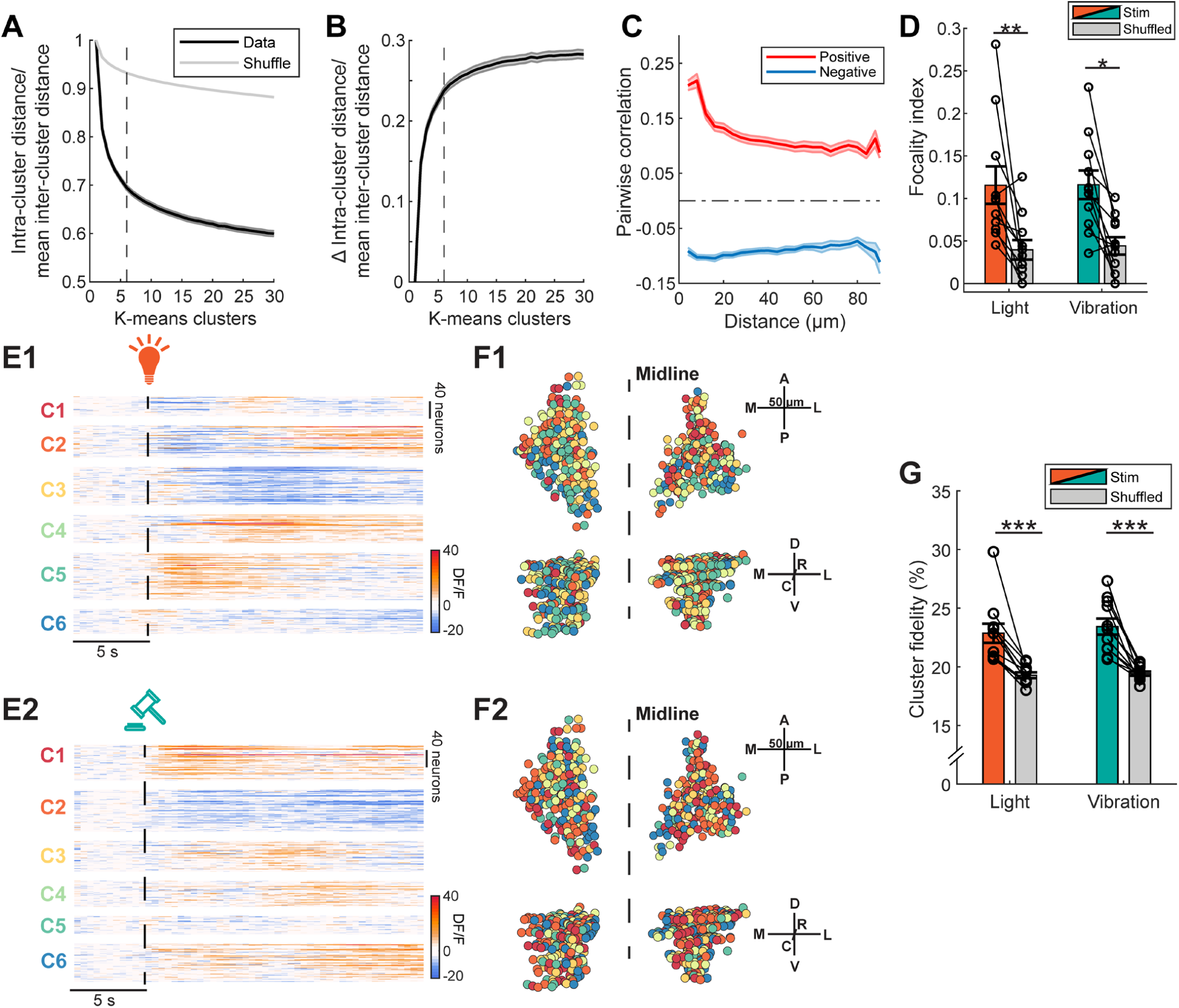
Preglomerular nucleus neurons form topographically organized functional ensembles with different sensory preferences. **A**. Identification of relevant and smallest number of k-means clusters to represent the ongoing spontaneous preglomerular nucleus activity, using the elbow method (see methods). The dashed line highlights the k-means analysis for 6 clusters to be a number that was the smallest possible and relevant, hence optimal. Shading represents the standard error of the mean. **B**. Difference between black and grey curves in A. Note that the relevant number of 6 clusters is near the region where the curves slope decreases. **C**. The positive (red) and negative (blue) pairwise correlations of all PG neurons during the spontaneous activity period were plotted as a function of distance between neuron pairs (N = 11 fish). **D**. The focality index measures the spatial distribution of the responding neurons in the PG where 1 indicates all responding neurons are located in a single focus, 0 means randomly distributed organization of sensory responses within PG. The focality index for all responding PG neurons during the light (left) and vibration (right) responding periods were compared to a control distribution where the spatial locations of each neuron were randomly shuffled (N = 11 fish, * p < 0.05, ** p < 0.01, Wilcoxon signed-rank test). **E**. Mean sensory response time courses for all PG neurons in an example fish following light (E1) and vibration (E2) stimulation. PG neuron responses were clustered using k-means clustering where the starting centroid was the same as in the spontaneous activity periods in Fig. 2. **F**. 3D reconstruction of the PG neurons color-coded using the cluster identities following light (F1) and vibration (F2) stimulations. **G**. The cluster fidelity is the fraction of neuron pairs remaining in the same k-means cluster during ongoing activity versus sensory response periods. The cluster fidelity of the light (left) and vibration (right) responding periods were compared to a control distribution where the cluster indices were shuffled (N = 11 fish, *** p < 0.001, Wilcoxon signed-rank test). All error bars are shown as standard error of the mean.

**fig. S5.**
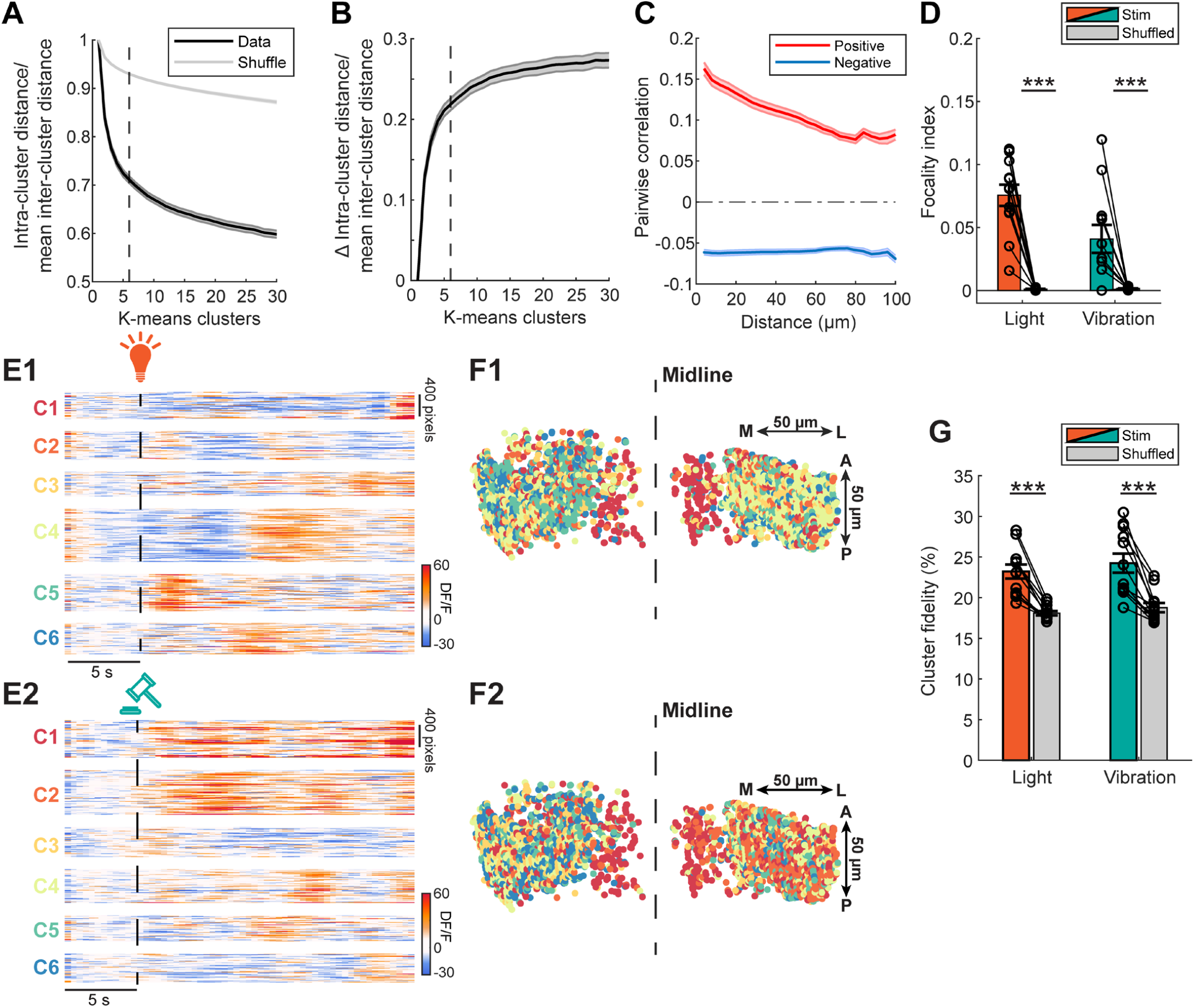
Preglomerular nucleus axons in the pallium form topographically organized functional ensembles with different sensory preferences. **A**. Identification of relevant and smallest number of k-means clusters to represent the ongoing spontaneous preglomerular nucleus axonal activity in the zebrafish pallium, using the elbow method. The dashed line shows the k-means analysis for 6 clusters to be optimal. Shading represents the standard error of the mean. **B**. Difference between black and grey curves in A. Note that the relevant number of 6 clusters is near the region where the curves slope decreases. **C**. The positive (red) and negative (blue) pairwise correlations of all PG axonal pixels during the spontaneous activity period were plotted as a function of distance between pixel pairs (N = 12 fish). **D**. The focality index for all responding PG axons during the light (left) and vibration (right) responding periods were compared to a control distribution where the spatial locations of each pixel unit were shuffled randomly (N = 12 fish, *** p < 0.01, Wilcoxon signed-rank test). **E**. Mean sensory response time courses for all pallial PG axons in an example fish following light (E1) and vibration (E2) stimulation. The PG axonal responses were clustered using k-means clustering where the starting centroid was the same as in the spontaneous activity periods in Fig. 4. **F**. 3D reconstruction of the pallial PG axons color-coded using the cluster identities following light (F1) and vibration (F2) stimulations. **G**. The cluster fidelity of the light (left) and vibration (right) responding periods were compared to a control distribution where the cluster indices were shuffled (N = 12 fish, *** p < 0.001, Wilcoxon signed-rank test). All error bars are shown as standard error of the mean.

**fig. S6.**
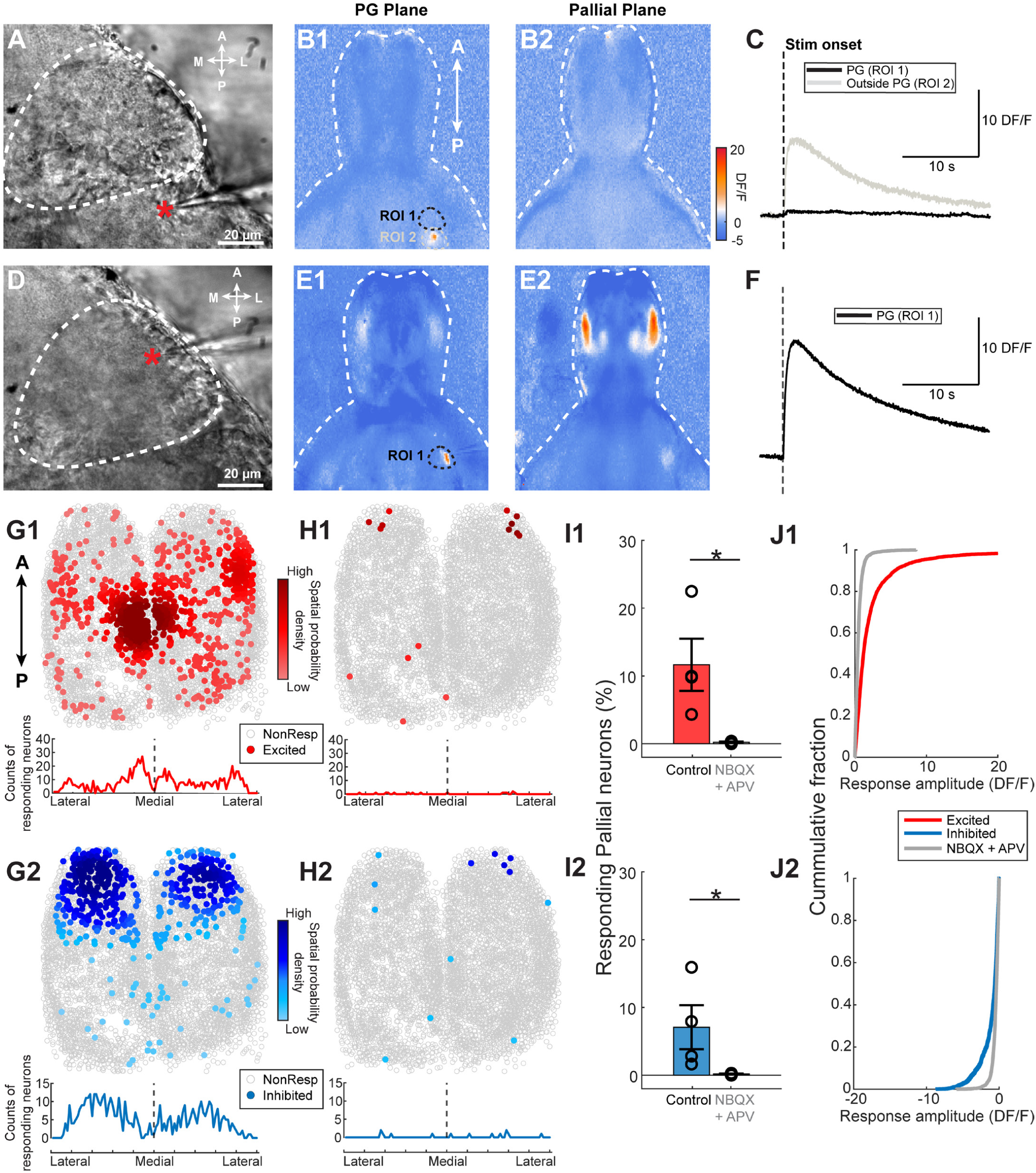
Local micro-electrode stimulation of preglomerular neurons evoked localized pallial responses, which are abolished with glutamatergic blockers. **A**. Brightfield image of a control micro-electrode stimulation experiment targeting an area (red star) adjacent to the PG (delineated by the white dashed line) in an example juvenile zebrafish brain explant. **B**. Baseline-subtracted epifluorescent (DF/F) image of the optical plane containing the PG (B1) and the pallium (B2). The white dashed line outlines the brain while the black region of interest (ROI 1) highlights the location of the PG and the gray ROI 2 highlights the region outside of the PG. **C**. Mean responding trace of the region of interest containing the PG (in black) and the ROI adjacent to the PG (in gray) in an example fish. **D**. Brightfield image of a micro-stimulation experiment targeting the PG (red star) which is delineated by the white dashed line in the same example fish as in A. **E**. Baseline-subtracted fluorescent (DF/F) image of the plane containing the PG highlighted by ROI 1 (E1) and the pallium (E2) following the stimulation in PG. **F**. Mean calcium trace of the PG following the focal micro-electrode stimulation in D. **G**. Top: 2D reconstruction of all excited (G1, red) and inhibited (G2, blue) pallial neurons and non-responding neurons (in gray) upon PG stimulation. The data from all fish are spatially aligned and overlaid. The color gradient represents the spatial probability density of the responding pallial neurons. Bottom: Number counts of excited (red) and inhibited (blue) neurons along the lateral-medial axis (N = 4 fish). **H**. Top: 2D reconstruction of the excited (H1) and inhibited (H2) pallial neurons of the same fish as in G following the bath application of glutamatergic receptor blockers (10 μM NBQX and 50 μM APV). Bottom: Number counts of excited (red) and inhibited (blue) neurons along the lateral-medial axis (N = 4 fish). **I**. The fraction of excited (I1) and inhibited (I2) neurons before and after the application of glutamatergic receptor blockers (N = 4 fish, * p < 0.05, Student’s t test). **J**. The cumulative fraction of the excited (J1) and inhibited (J2) neurons before (colored) and after (gray) the application of synaptic blockers. Error bars denote the standard error of the mean (N = 4 fish).

**fig. S7.**
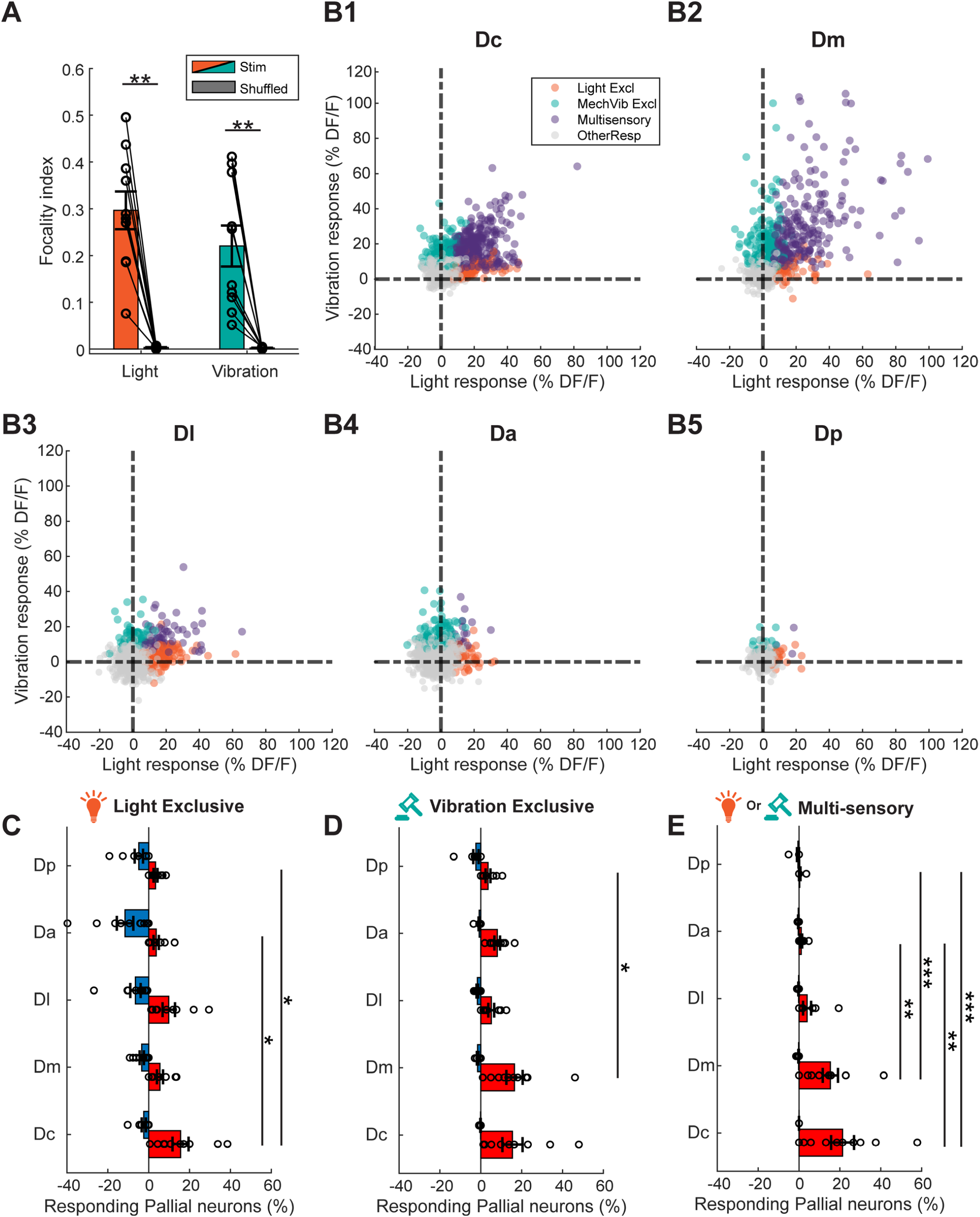
Sensory response preferences across distinct pallial brain regions are heterogenous. **A**. The focality index for all responding pallial neurons following light (left) and vibration (right) stimulations were compared to a control distribution where the distances between neurons were shuffled. (N = 10 fish, ** p < 0.01, Wilcoxon signed-rank test). **B**. Scatter plot of the mean sensory responses of all neurons in anatomically delineated pallial regions: Dc (B1), Dm, (B2), Dl (B3), Da (B4), Dp (B5). Neurons with significant excitation to at least 1 of the sensory stimuli are color-coded based on their response selectivity for either light (orange), vibration (teal) or both (purple). All other neurons are shown in gray. **C-E**. Fraction of responding pallial neurons in the delineated brain regions according to their sensory selectivity: light exclusive (C), vibration exclusive (D), or multi-sensory (E). Red denotes excited and blue denotes inhibited pallial neurons following sensory stimulation (N = 10 fish, * p < 0.05, ** p < 0.01, *** p < 0.001, Kruskal-Wallis test followed by Dunn’s test). Error bars denote the standard error the mean.

**fig. S8.**
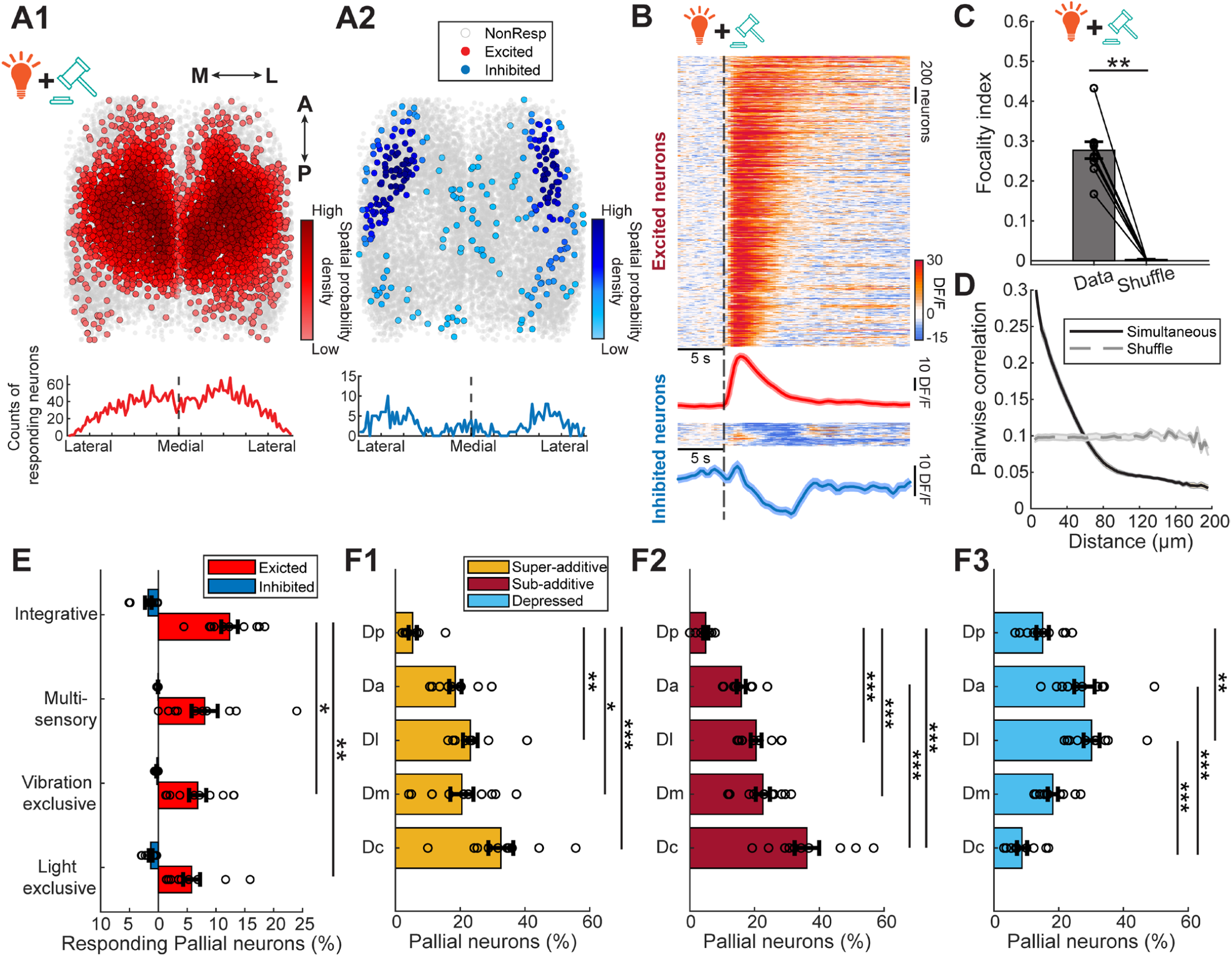
Integrative pallial neurons that are activated by the simultaneous delivery of sensory stimuli are topographically organized in the zebrafish pallium. **A**. Overlaid 2D reconstruction of the excited (in red, A1) and inhibited (in blue, A2) pallial neurons from all spatially aligned fish, in response to the simultaneous presentation of both light and vibration stimulation, in vivo. The color gradient represents the spatial probability density of the responding pallial neurons. Non-responding pallial neurons are in gray. **B**. Time courses of the pallial neurons’ calcium signals in response to the simultaneous presentation of both light and vibration stimuli. Warm colors indicate increased neural activity; cold colors indicate decreased activity. Mean calcium signals are shown at the bottom of each heatmap; shades indicate the standard error of the mean (N = 10 fish) **C**. The focality index for all responding pallial neurons during the light (left) and vibration (right) responding periods were compared to a control distribution where the spatial locations of each neuron were shuffled randomly (N = 10 fish, ** p < 0.01, Wilcoxon signed-rank test). **D**. The pairwise correlation of the pallial neurons’ responses to the simultaneous delivery of sensory stimuli as a function of the distance between neuron pairs. The grey dashed line represents the pairwise correlations when the distances are shuffled. The shaded region denotes the standard error of the mean. (N = 10 fish). **E**. Fraction of responding pallial neurons according to their sensory response selectivity; light exclusive, vibration exclusive, multi-sensory and integrative. Excited neurons are in red and inhibited neurons in blue (N = 10 fish, * p < 0.05, ** p < 0.01, Wilcoxon rank-sum test). **F**. Fraction of pallial neurons in anatomically delineated pallial regions based on their sensory response characteristics; super-additive (F1), sub-additive (F2), or depressed (F3) (N = 10 fish, * p < 0.05, ** p < 0.01, *** p < 0.001, Kruskal-Wallis test followed by Dunn’s test). All error bars denote the standard error of the mean.

